# Patient-Specific Gene Co-Expression Networks Reveal Novel Subtypes and Predictive Biomarkers in Lung Adenocarcinoma

**DOI:** 10.1101/2024.08.06.606802

**Authors:** Patricio Lopez-Sanchez, Federico Avila-Moreno, Enrqiue Hernandez-Lemus, Marieke L. Kuijjer, Jesus Espinal-Enriquez

## Abstract

Lung adenocarcinoma (LUAD) is a highly heterogenous and aggressive form of non-small cell lung cancer (NSCLC). The use of genome-wide gene co-expression networks (GCNs) has been paramount to describe changes in the transcriptional regulatory programs found between diseased and healthy states of LUAD. Recently, studies have shown that multiple cancerous phenotypes share a distinct GCN architecture, suggesting that network topology holds promise for understanding disease pathology. However, conventional GCN inference methods struggle to capture the inherent context-specificity within a patient population, thus flattening its heterogeneity. To address this issue, the use of single-sample network (SSN) modelling has emerged as a promising solution into studying heterogeneous traits of cancer through network-based approaches. Here, we reconstructed patient-specific GCNs (n=334) using the LIONESS equation and mutual information as the network inference method. Unsupervised analysis revealed six novel LUAD subtypes based on inter-patient network similarity, each with distinct network motifs reflecting unique biological programs. Supervised analysis, employing regularized Cox regression, identified 12 genes (CHRDL2, SPP2, VAC14, IRF5, GUCY1B1, NCS1, RRM2B, EIF5A2, CCDC62, CTCFL, XG, andTP53INP2) whose weighted degree in SSNs is predictive of patient survival in LUAD. These findings suggest that topological features of SSNs offer valuable insights into the context-specific nature of LUAD malignancy, highlighting the potential of SSN-based approaches for further research.

## 1 Introduction

Lung cancer remains the most fatal and second most frequently diagnosed cancer globally. In 2020, it constituted 18% of cancer-related deaths and was responsible for 11.4% of new cancer cases[1].

Lung cancer can be classified into two main sub-types: non-small cell lung cancer (NSCLC) and small cell lung cancer (SCLC), with NSCLC accounting for 85% of all newly diagnosed cases. Adenocarcinomas and squamous-cell carcinomas are the two most common subtypes of NSCLC, constributing to 50% and 30% of all NSCLC cases, respectively [2]. Among NSCLCs, squamous cell carcinomas often arise in the central bronchial tubes and express molecular markers of squamous cell differentiation. In contrast, adenocarcinomas typically develop in the lungs’ periphery and are frequently associated with the formation of glandular bodies and high mucin production. Both cancer subtypes exhibit significant differences in clinical associations such as gender, age, smoking history, TNM clinical stages, 5-year survival rates post-diagnosis, and treatment options, among others [3].

Lung cancer cells exhibit aberrant regulatory programs at multiple physiological scales, ranging from their mechanisms of genetic and epigenetic regulation to the interaction between the tumor tissue and its surrounding environment[4]. For instance, genetic alterations in a set of proto-oncogenes such as KRAS, EGFR, BRAF, PI3K, MEK, and HER2 have been identified to severely dysregulate various signaling pathways crucial for the control of cell proliferation, apoptosis, and other cellular functions[5]. The disruption of these mechanisms directly influences the phenotypic transition from healthy to cancerous tissue. Therefore, there is a growing motivation to study the molecular biology of lung cancer, employing a wide variety of multiand interdisciplinary approaches with the aim of developing new strategies for prevention, early diagnosis, and treatment.

Within lung cancer, each subtype produces a large number of distinct pathological signatures, characterized by the differential interactions between the components in the system. To study these complex diseases, systems biology has sought to model these interactions through network-theoretical approaches. An example of this, and the main focus of this work, is gene co-expression networks (GCNs). GCNs are undirected graphs where nodes represent genes, and edges represent a statistically significant dependence between their expression levels[6]. The importance of GCN analysis in biomedicine lies in its ability to model interactions among genes expressed in a tissue, thus illustrating the landscape of their transcriptional program in healthy and diseased states.

Recent studies have revealed clear underlying topological differences between networks constructed from samples of cancerous tumors and those from adjacent healthy tissue. For example, a clear rupture of the largest connected component into smaller sub-graphs has been observed. Furthermore, comparative analyses of these networks has shown a clear reduction in the links between genes located on different chromosomes, while interactions between genes physically close to each other on the genome are greatly enhanced. This phenomenon, coined as the loss of longdistance co-expression, has been observed in various malignant tumors, including Luminal A, Luminal B, HER2-positive, and basal breast cancer[7] [8], clear cell renal carcinomas[9], NSCLCs [10], and more recently in hematological cancers[11], thus suggesting that this is a common trait of cancer. These results highlight that genome-wide GCN topology is intrinsically associated with cancer development. However, understanding how global or local network structural traits can be linked to the underlying disease’s biology remains a topic of wide discussion.

Given that GCNs capture statistical dependence between genes, network inference algorithms require multiple samples to construct a robust aggregate network that accurately represents a sample population or cohort. One of the major limitations of these methods is that they can only represent characteristics shared by the chosen set of background samples, therefore losing relevant information about the specific context and environment of each sample in the process. Thus, while extremely useful at mining shared traits in a biological phenomenon, aggregate networks are unable to capture the underlying heterogeneity that exists within the population of interest.

To better understand how phenotypic complexity varies across individual samples from a network-driven perspective, the LIONESS method (Linear Interpolation to Obtain Network Estimates for Single Samples) for estimating sample-specific networks (SSNs) was developed[12]. This method is based on the assumption that aggregate networks are the result of a linear combination of individual sample networks. As a first step, the LIONESS equation contrasts two input networks: one built on data from all samples and another excluding data from a single sample. This step essentially calculates the contribution of the removed sample to the aggregate network. The estimated contribution is then normalized to account for sample size and added to the aggregate network without the sample of interest, effectively estimating that sample’s individual network. This process can then be repeated iteratively to infer a unique network for each sample within the population. The relevance of LIONESS lies in its potential to reveal connections between GCNs’ topological features and clinically relevant factors, such as overall survival. By enabling analysis at the individual sample level, LIONESS-based single sample GCNs could offer a deeper understanding of how variations in gene-correlation patterns might influence disease progression and patient outcomes.

Here, by using LIONESS, we have inferred patient-specific GCNs of the LUAD-TCGA cohort using mutual information as the network reconstruction algorithm, and subsequently developed two parallel frameworks for their analyses. The first approach consisted of a graph-clustering strategy whereby focusing on communities of patients based of their on SSN-similarity, we explored the significant relationship between conserved network motifs and LUAD clinical, molecular and cellular contexts. The second one, is focused in node weighted degrees; we developed a regularized regression scheme to extract genes whose structural importance in a SSN is predictive of LUAD overall survival. We then used random resampling methods in order to ascertain the robustness of our findings.

## 2 Methods

### 2.1 RNA-Sequencing Data Acquisition and Preprocessing

Following the methodology established by Andonegui et al [10], clinical and RNA-sequencing (RNA-Seq) data corresponding to patients within the Lung Adenocarcinoma (LUAD) project in The Cancer Genome Atlas (TCGA) Research Network: https://www.cancer.gov/tcga were extracted using the TCGAbiolinks R package [13]. Subsequently, all transcripts that were not protein-coding genes were filtered out, and genes with a zero-prevalence higher than 50% across all samples were removed. The resulting expression matrix was normalized to correct for transcript length effects, GC content, and library size. Furthermore, in order to focus on overall patient survival and explore its association with SSNs’ topological features, samples belonging to patients whose survival data were not properly annotated were excluded. Thus, the analysis centered solely on patients who had been followed for more than a year or who had died before this period elapsed. Next, the scattering of expression profiles was visualized using principal component analysis (PCA) to verify the proper preprocessing of the samples. Finally, in order to focus on LUAD disease heterogeneity, only samples from primary tumor tissue were kept for SSN building and downstream analyses.

### 2.2 Single sample network reconstruction and modelling

The LIONESS equation requires two objects as input: the first is a GCN built using all samples (network *a*) and the second is a GCN built without sample *q* (network *a* − *q*). To build the two input GCNs, we use ARACNE’s implementation of Mutual Information (MI) as the measurement of the statistical dependence between the expression of all pair-wise combinations of genes [14]. In order to handle the inputs for both ARACNE and LIONESS in a computationally efficient manner, we integrated LIONESS into a multi-core backend for ARACNE [10] so that all MI calculations are done in parallel, while each SSN is built sequentially.

By using this workflow, we were able infer a total of 334 patient-specific LUAD GCNs. To allow the individual inspection of each network’s nodeand edgelevel topological description, we created fully-connected SSNs as well as subsets of the top 10,000, 50,000 and 100,000 edges ranked by their weight (i.e. LIONESS score).

### 2.3 Clustering of patient network-similarity matrix

To assess whether the higher order structure of Individual GCNs is shared among phenotypically similar patients, we constructed a Patient network-similarity matrix *A*, where each entry *a_ij_* contains the Jaccard-similarity index between the sets of edges of the filtered networks *i* and *j*. *A* is an *n* × *n* symmetrical matrix where *n* is the number of SSNs, making it suitable for representation as an adjacency matrix of an undirected weighted graph. In this graph, nodes represent patients and edges represent their SSN similarity score. This notation enables us to reframe the problem of patient clustering based on SSN similarity as one of detecting community structures in a graph. We performed community detection using the Louvain algorithm for weighted networks as implemented in the igraph R package [15] [16]. Varying resolution values were allowed for the detection of smaller communities, as it is known that graph clustering methods based on modularity maximization can be biased towards detecting larger clusters in a network[17].

### 2.4 Biological assessment of patient network-similarity clusters

Communities obtained from the Patient-similarity-graph were probed to shed light into the traits that drive clinical, cellular and molecular phenotypic heterogeneity. Thus, to biologically interpret each patient-cluster, we seek to answer three questions:

1.-Are patients in a given cluster significantly enriched for particular clinical features?
2.- Do patient-clusters have distinct cellular compositions?
3.- What are the main biological processes that drive the formation of heterogenous phenotypes in the population?

To address the first question, we used hypergeometric testing followed by FDR correction to investigate statistically significant (q-value *<* 0.05) over-representation of gender, disease stage, tumor size and extent (T), lymph node involvement (N), and presence of distant metastasis (M). Moreover, we used a log-rank test to look for significant differences (p-value *<* 0.05) in the survival distributions of patients between clusters.

For the second question, we utilize the R implementation of Xcell [18] to calculate cell enrichment scores for each sample using normalized expression values to account for transcript length. We then conduct Kruskal-Wallis tests with post-hoc Dunn’s test to identify any statistically significant associations between different cell types’ enrichment score (Xcell value) and patient communities.

Lastly, to extract the most representative biological processes in each community, we repurposed the CoDiNA (Co-Expression Differential Network Analysis) framework to identify the most conserved edges within each cluster of patient-specific networks [19]. In essence, we obtain the entire edge-space of a given cluster of networks and examine the frequency with which an edge appears throughout all samples. We then consider an edge to be highly conserved in a set of networks if it is within the 99th percentile of most frequent edges. Thus, by aggregating the most conserved interactions, we are able to construct consensus networks that portray the most active set of sub-graphs in each cluster.

Subsequent biological annotation of each per-cluster consensus network is achieved by querying the presence of significantly enriched (q-value *<* 0.01) Biological Processes in its largest connected component. The detection of the largest connected components and functional enrichment analysis were carried out using the clusterMaker [20] and gProfiler [21] Cytoscape plugins [22], respectively.

### 2.5 Calculation of genes’ weighted degrees in Single-sample Networks

The interaction between genes are severely altered during the transition from healthy to diseased states. These alterations can be better summarised by examining the centrality of individual nodes in each network. For example, several studies have previously used the weighted in-degrees of patient-specific regulatory networks as proxy of the amount of regulation that a given gene receives from all transcription factors [23] [24] [25]. Drawing inspiration from this approach, we built an *m*×*n* matrix of weighted degrees (WDs) for all *m* protein coding genes in *n* fully-connected SSNs. In the context of individual GCNs, the weighted degree of a gene serves as a summary statistic indicating the extent of potential active interactions within a SSN, encompassing both direct and indirect regulation, as well as patterns of coregulatory behaviour. Weighted degrees for all genes in each SSN were calculated using the igraph R package.

To validate that the resulting WD matrix reflects true biology of LUAD, we performed pre-ranked gene set enrichment analysis (GSEA; [26] [27]) on the standard deviation values of all WD scores. This allows us to investigate whether genes with the most variable patterns of connectivity are associated to specific Biological Processes [28] or Hallmark gene sets[29]. We use FDR *<* 0.01 as our significance cutoff in each enrichment test.

### 2.6 Predictive modelling of patient overall survival through penalized regression

To identify the sets of genes whose weighted degree in a network is associated to survival in patients with LUAD, we utilized the WD matrix as features for a penalized Cox proportional hazards regression model while correcting for gender and age at diagnosis [30]. Model construction was performed on a 70% training split, while validation was carried out on the remaining 30% hold-out test data set. In particular, given the high dimensionality of the WD matrix, we make use of the L1 penalty as it is known that this type of regularization performs both feature selection and model fitting in a manner that favours sparser models and counters overfitting [31] [32].

We built two different regression models using the Least Absolute Shrinkage and Selection Operator (Lasso) penalty. The first involves using the complete WD matrix when constructing the model, i.e. we let Lasso have full control over the selection of genes that are relevant to the final fit. The second model involves a univariate Cox proportional hazards screening step for each gene prior to the shrinkage procedure. Thus, only genes with a significant univariate association to survival (log-rank test, p-value *<* 0.05) were kept for subsequent Lasso selection. The motivation behind this double filtering step is to first reduce the amount of competing noisy variables that could bias the shrinkage of coefficients, followed by a second reduction of the model’s complexity by filtering out correlated variables through the L1 penalty.

Despite Lasso’s effectiveness in dealing with high dimensional data, it has been reported that Lasso tends to over shrink true large coefficients, thereby yielding biased estimates that are not directly suitable for generalization on test data [33]. To address this issue, the Relaxed Lasso has been proposed as a de-biasing procedure in which Lasso is only used as a variable screening step rather than a model selector. Consequently, the active set of variables are fitted through the Ordinary Least Squares solution without any penalization [34] [33]. Owing to this, we inspect if relaxing the models after Lasso variable selection produces a better fit. We utilized the relax.glmnet() function from the glmnet R package [35] [36] with gamma = 0 for model predictions and evaluation, as this produces the most relaxed fit possible.

Fitting a regularized model requires the fine-tuning of the penalization parameter lambda. To achieve this, we find the optimal values of lambda for each model variant through 10-fold cross validation. In particular, we choose the value of lambda that is one standard deviation away from the one that maximizes Harrel’s Concordance Index (C-Index;[37]) in the cross-validation procedure.

### 2.7 Model validation and feature stability analysis

To obtain a comprehensive evaluation of each model, we sought to quantify and compare all regressions’ goodness-of-fit, predictive performance and overall uncertainty.

As a goodness-of-fit test, and as a first illustration of each model’s prognostic power, we predicted risk scores for patients in the hold-out test data, and dichotomized them into High- and Low-risk groups using the median value as a threshold. Subsequently, we performed a log-rank test to check for significant differences between both risk groups’ survival distributions.

Next, to evaluate a survival model’s overall predictive performance, we focus on both global and time-discrete evaluation metrics. For a global assessment we utilize the C-Index as well as the Integrated Time-dependent Brier’s Score (IBS). To evaluate a model’s performance at discrete time points, we use the area under the receiver operator characteristic curve (AUROC) for 1-, 2-, 3- and 5-year overall survival predictions as well as the Time-dependent Brier’s Score (BS) for all possible time points.

Finally, to test each model-building procedure and measure the final overall performance’s uncertainty, we created 100 different training/validation data splits and repeated all filtering and hyperparameter-tuning steps, assessing the global discriminatory power in each run. Furthermore, we utilized this resampling-based method to identify the sets of genes most frequently selected by Lasso and the univariate filter. The frequency with which a gene was selected was then used as a representative measurement of the robustness of its association to patient survival.

## 3 Results

### 3.1 Most patient clusters based on single-sample network similarity are not associated with clinical phenotypes

As a first exploration of the extent to which SSN topology is capable of defining heterogeneous LUAD phenotypes, we investigated whether patients sharing a substantial number of their most important edges also shared similar clinical outcomes. **Figure 2** shows the presence of the community structure inside the SSN-similarity graph . Notably, Cluster 1 displays the subset of SSNs that have the highest resemblance to each other. On the other hand, Cluster 4 follows the opposite trend, where networks exhibit low similarity not only with those outside their community but also among themselves. To ensure that the observed pattern of community structure was not dependent on the selected threshold of 10,000 edges, we performed the same clustering strategy on sets of SSNs filtered to their top 50,000 and 100,000 edges. **Supplementary Figure 1** shows the overlap between the detected communities of SSNs at different network densities. As expected, clusters that were highly conserved such as clusters 1,2,3,5 and 6 were concordant with communities detected at varying SSN densities. Contrarily, cluster 4 was prone to be sub-clustered when considering additional interactions.

**Fig. 1.**
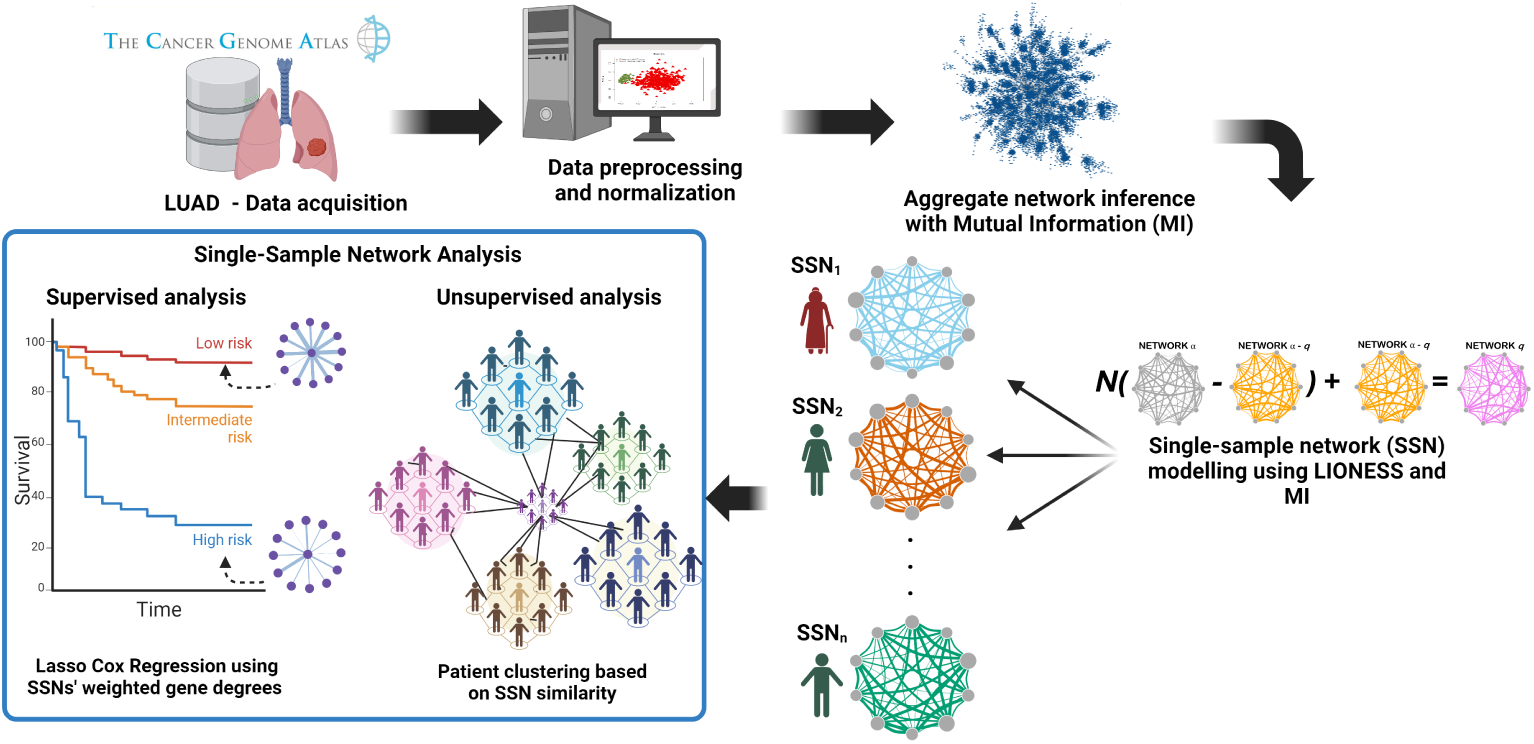
Schematic overview of the methods used in this study

**Fig. 2.**
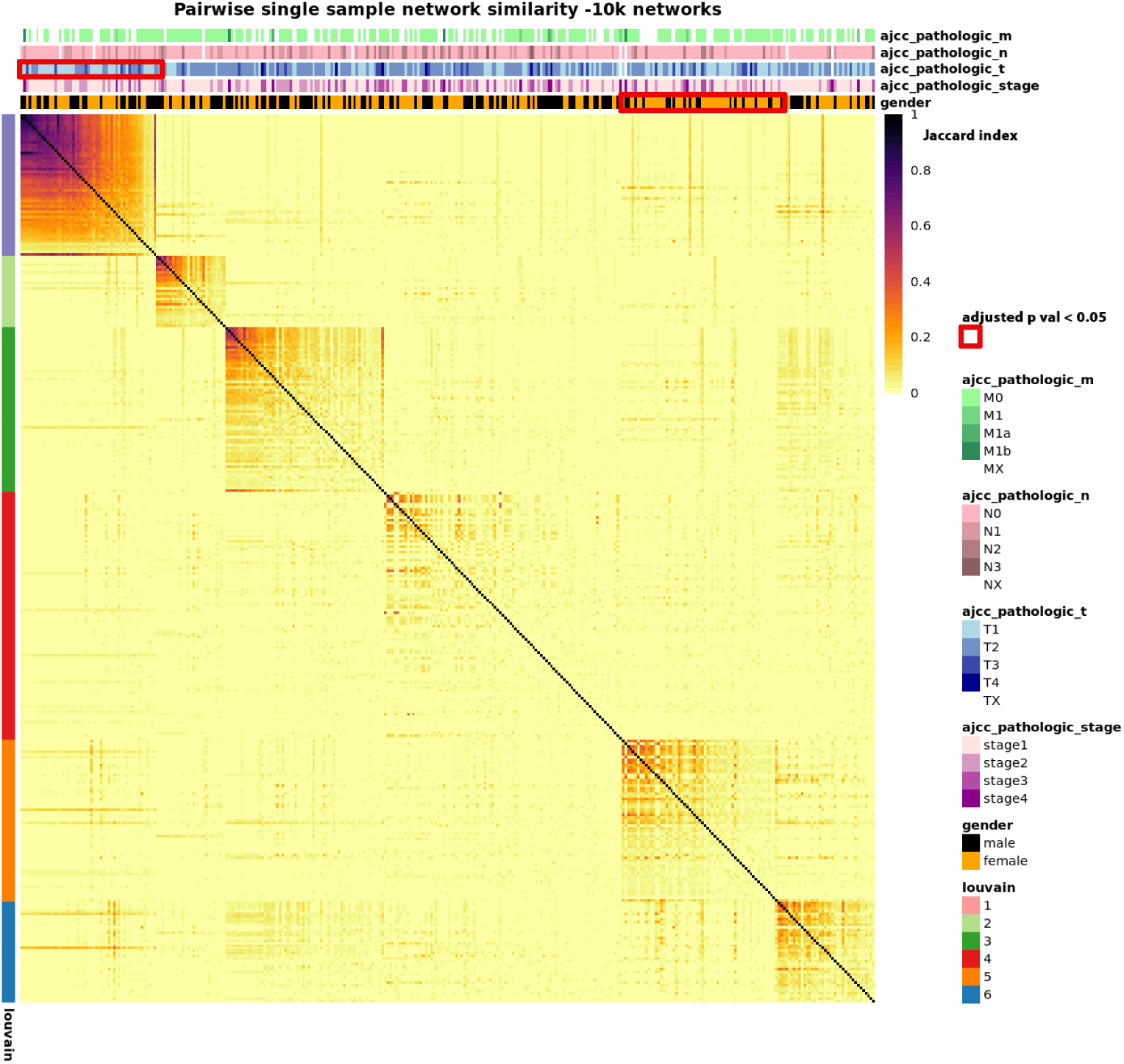
Similarity matrix between patient-specific networks. The top 10,000 edges of each network were used to calculate all pairwise Jaccard indices. The resulting weighted adjacency matrix was clustered using the Louvain algorithm. Row annotations show the clustering results. Column annotations show the different clinical labels associated to each patient. Red marques show the significant enrichment (Fisher’s exact test adjusted p-value *<* 0.05) of a clinical feature inside a cluster.

Next, we conducted overrepresentation testing within patient clusters to determine if clinical outcomes could be discerned from network similarity alone. Factors considered included gender, disease stage, tumor size and extent (T), lymph node involvement (N), and presence of distant metastasis (M). Interestingly, only Cluster 1 and Cluster 5 showed significant enrichment (Fisher’s exact test, adjusted p-value *<* 0.05) with T1 tumors and female patients, respectively (red marques, **Figure 2**). In addition, when comparing the survival distributions between clusters, no significant differences (log-rank test p-value *<* 0.05) were observed.

### 3.2 Per-cluster consensus networks are associated with cell and tissue context

To further characterize the biology behind the formation of patient communities based on SSN similarity, we built six consensus networks (one for each group) by identifying the most conserved set of interactions in each case (see Methods). Given that a percentile based statistic was used to determine if an interaction is conserved, consensus network size and density largely depend on the range of possible edges within each cluster. **Supplementary Table 1** contains the structural summary statistics of each consensus network.

It can be observed that the number of nodes, edges and connected components varies accordingly with the degree of similarity shared between SSNs in a cluster, hampering downstream comparative analyses . In consequence, to facilitate the annotation of all consensus networks, we focused solely on each largest connected component to perform subsequent functional enrichment analysis (**Figure 3**).

**Fig. 3.**
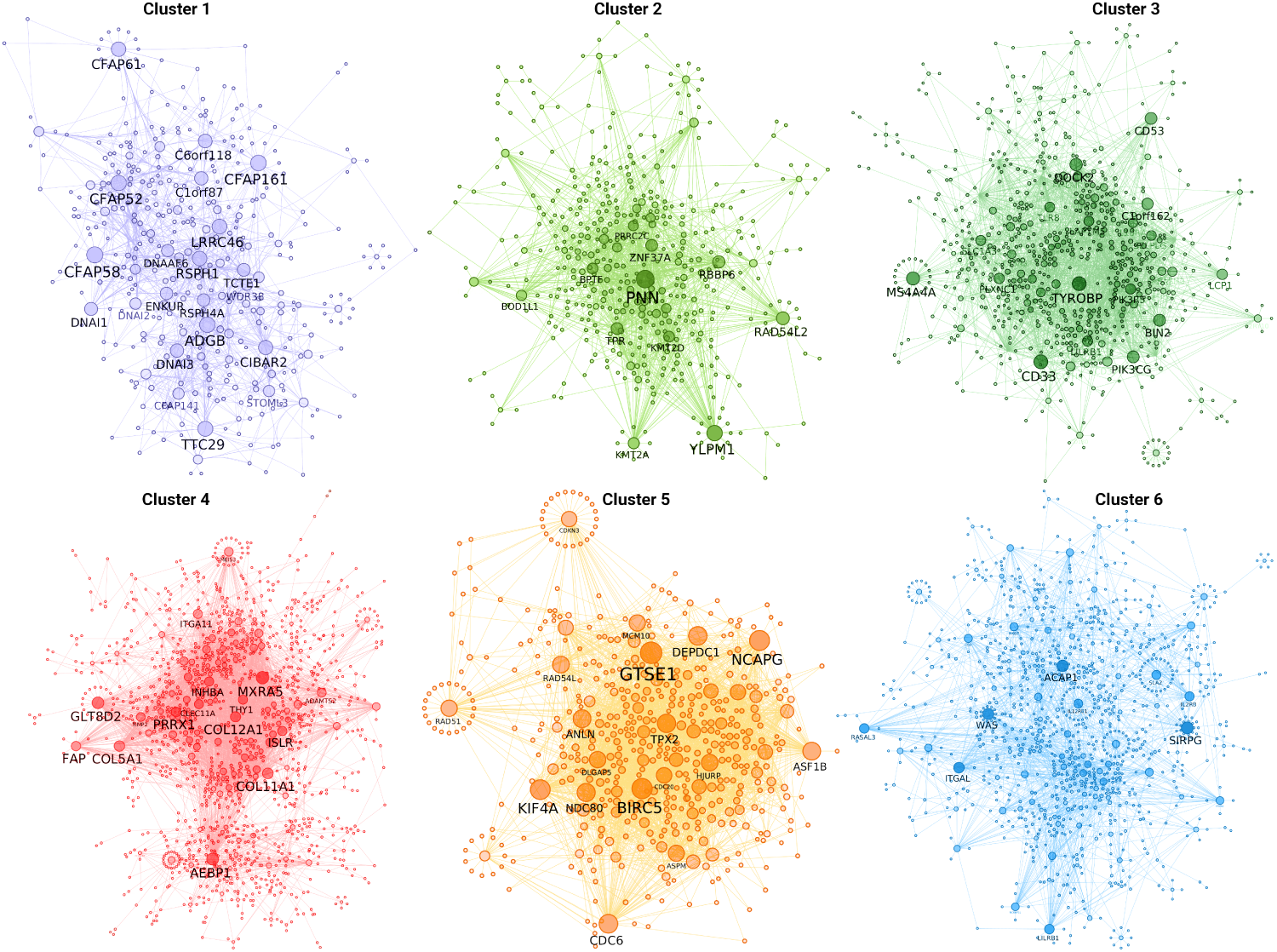
Largest connected components of per-cluster consensus networks. The top 1% most frequent edges in each cluster were aggregated to build the corresponding consensus network. Node and label size correspond to the number of incident edges (degree).

**Figure 4** depicts the biological processes significantly associated with each cluster (Fisher’s exact test, adjusted p-value *<* 0.05). It is possible to note that, with the exception of Cluster 3 and Cluster 6, each patient community is enriched by unique biological programs: Cluster 1 is associated with processes related with cilium movement and structure. Cluster 2 is associated with processes regarding chromatin remodelling and histone modification. Cluster 4 is enriched with angiogenic and matrix remodelling processes. Cluster 5 is strongly associated with cell cycle progression and regulation. Finally, Cluster 3 and Cluster 6 share multiple enriched processes related to the activation and regulation of the immune response.

**Fig. 4.**
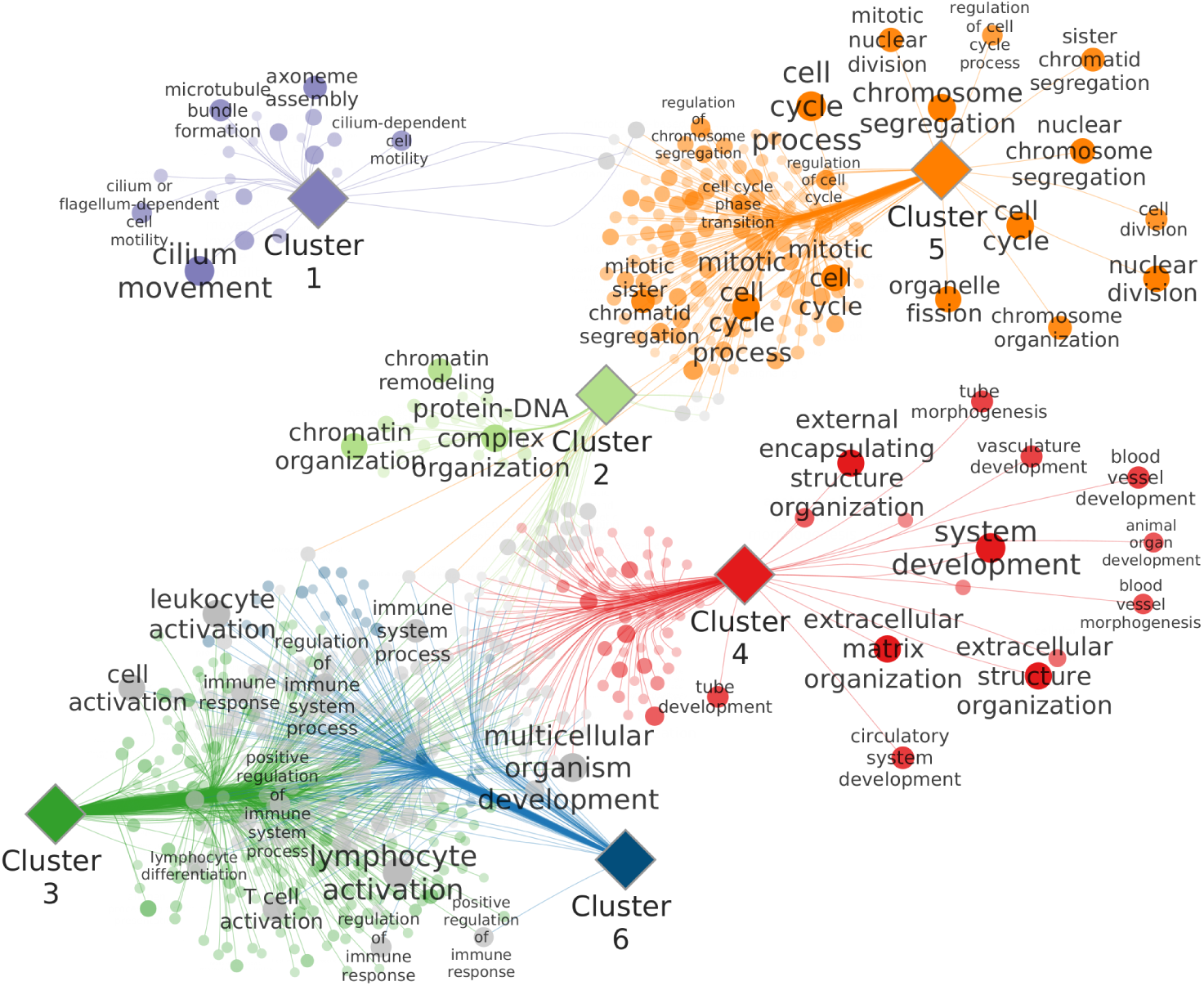
Bipartite network of enriched Biological Processes in each consensus network. Diamond-shaped nodes represent different clusters of patients, while circular-shaped nodes represent a Biological Process enriched in its corresponding network’s largest connected component. Node and font size and transparency vary according to the scaled -log(adjusted p-value) in the enrichment test. Grey colored nodes indicate Biological Processes that are enriched in more than one Cluster, while nodes with the same color as their neighbor Cluster node represent a Biological Process specific to that group.

Considering that, despite having dissimilar networks, Cluster 3 and Cluster 6 shared the vast majority of overrepresented immune system-related biological processes, we sought to evaluate whether this phenomenon could be caused by underlying differences in tumor cell composition. We used the R implementation of xCell to estimate cell enrichment scores for 64 immune and stromal cell types from expression data. We identified 18 immune cell types and 11 stromal cell types with significantly different distributions (Kruskal-Wallis H, adjusted p-value *<* 0.001) in at least one patient cluster (**Figure 5A**). A post-hoc Dunn test of xCell’s Immune Score shows that, in general, the immune cell composition of Cluster 3 and Cluster 6 is inversely correlated (**Figure 5B**), with Cluster 3 being the most immune-depleted group of patients, while Cluster 6 displays the most infiltrated group of tumors. Analogously, the Stromal Score distribution of Cluster 3 follows a similar trend by displaying the lowest degree of infiltration (**Figure 5C**). Overall, as a summary for both stromal and immune cell types, xCell’s Microenvironment Score exhibits the same pattern as the one shown in immune cell types (**Figure 5D**), suggesting that larger inter-cluster changes in microenvironment composition are mainly driven by the immune milieu.

**Fig. 5.**
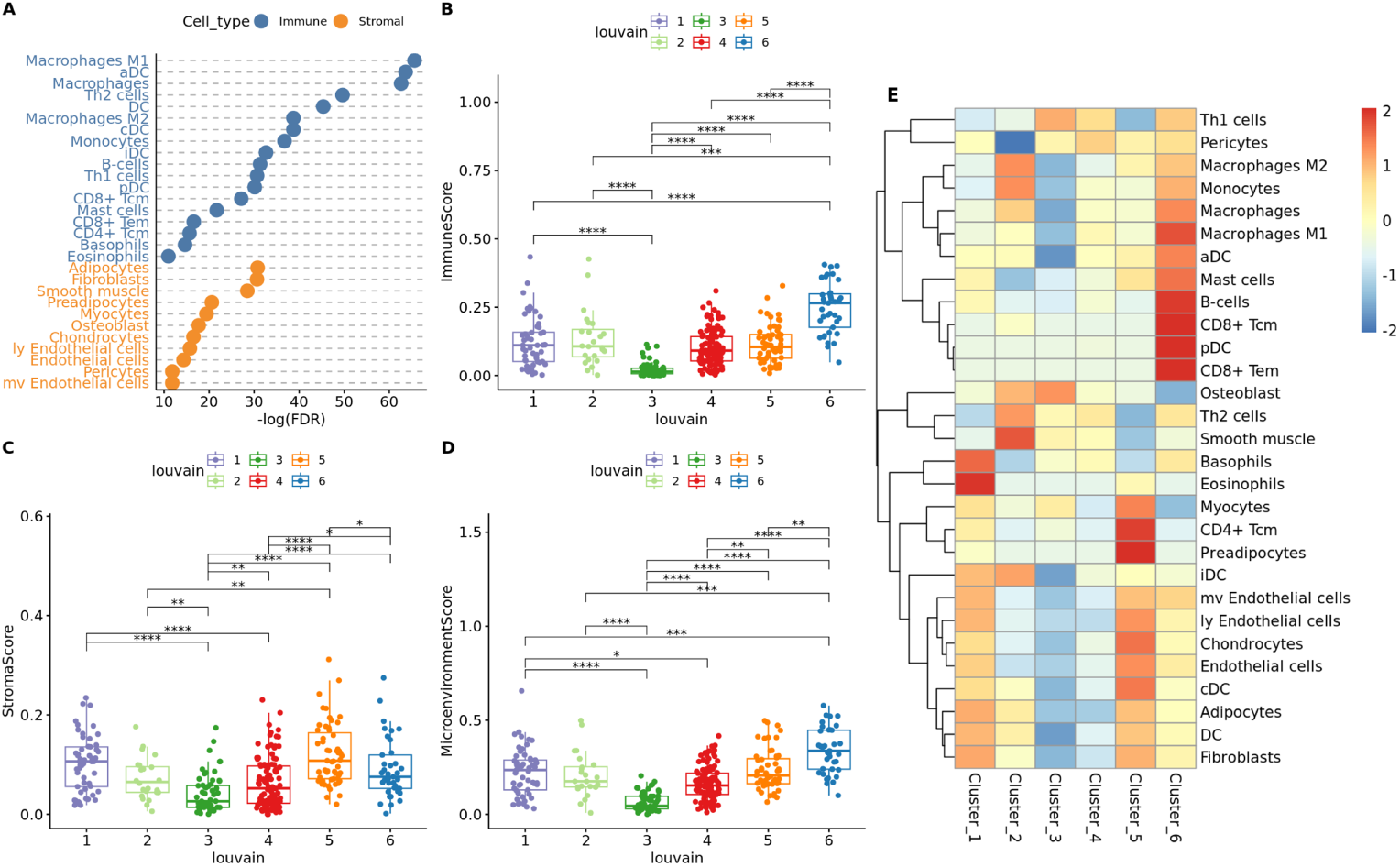
Differences in cell enrichment scores across clusters. **A:** Dot-chart of Kruskal-Wallis test results comparing xCell’s immune and stromal cell type enrichment scores between patient communities. Significance threshold was set at adjusted p-value *<* 0.001. **B-D:**Distribution of estimated ImmuneScore, StromalScore and MicroenvironmentScore, respectively. Pairwise significant differences were calculated through post-hoc Dunn testing. **E:** Heatmap of the scaled median Xcell values for all cell types with significant differences obtained in the Kruskal-Wallis test.

Finally, **Figure 5E** illustrates the variation in median xCell values for all significant cell types across different patient communities. From a more localized perspective, each cluster exhibits a distinct cell infiltrate signature involving both immune and stromal cells: Cluster 1 and Cluster 5 are the groups with highest enrichment of multiple endothelial cell types. However, Cluster 1 displays the highest enrichment of Neutrophils and Basophils while Cluster 5 shows the strongest signal of CD4+ Tcm and Preadipocytes. In parallel, Cluster 2 is characterized by the hightest levels of Th2 cells, M2 macrophages, iDCs (immature dendritic cells) and smooth muscle cells, as well as a depletion of pericytes. Cluster 3 cell enrichment signature is consistent with what is shown is the three summary scores for immune, stromal, and microenvironment cell compositions, displaying a depletion of most cell types, with the notable exception of high levels of Osteoblasts and Th1 cells. Cluster 4 displays an overall stable signature of all highlighted cell types, while Cluster 6 displays the strongest signal of immune infiltration, characterized by M1 macrophages, aDCs (activated dendritic cells), pDCs (plasmacytoid dendritc cells), B-cells, CD8+ Tcm (T central memory) cells and CD8+ Tem (T effector memory) cells.

### 3.3 Weighted Gene Degrees capture structural changes associated with cell proliferation

As a way of summarizing the role that an individual gene plays in the overall structure of each SSN, we calculated the weighted degree of each gene as the sum of the weights of all incident edges. With the goal of investigating whether specific hallmarks or biological processes rewire their connectivity the most, we conducted gene set enrichment analysis on the standard deviation value of each gene’s weighted degree. Noticeably, 5 hallmarks associated with cell proliferation, (MYC Targets V1, MYC Targets V2, E2F Targets, G2M Checkpoint and Mitotic Spindle) are among the categories with highest enrichment scores (**Figure 6A**). Comparably, when looking for enriched Biological Processes, we observed a similar pattern where multiple programs related to cell proliferation, were significantly enriched (**Figure 6B**). These results suggest that the network of genes regulating cell cycle progression and replication is under constant structural change. Additionally, weighted gene degrees, as a topological feature of GCNs, reflect biological phenomena and provide a stable foundation for downstream analyses.

**Fig. 6.**
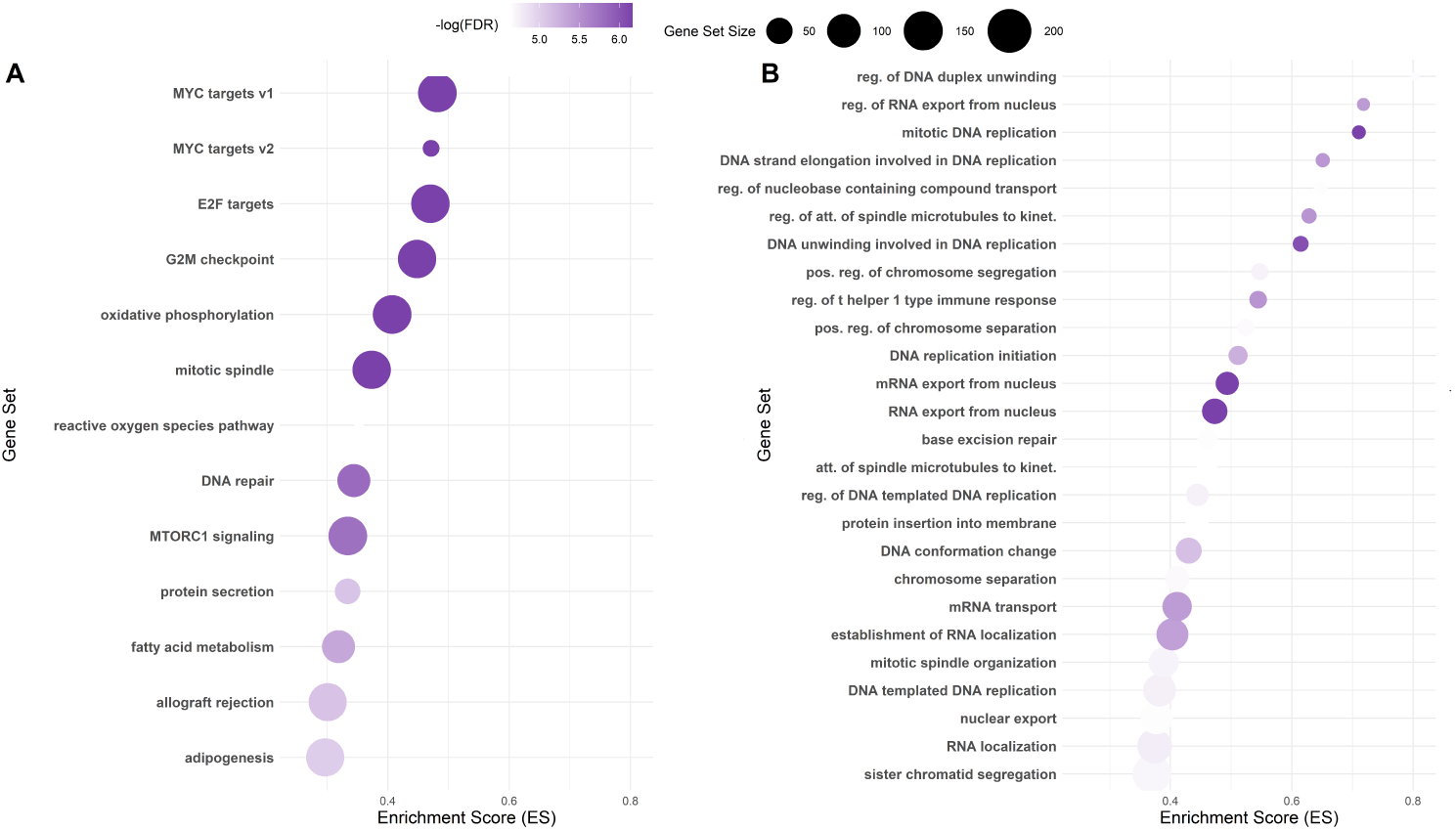
Gene Set Enrichment Analysis (GSEA) results of significant (FDR *<* 0.01) **A:**Hallmark signatures, and **B:** Biological Processes, according to the standard deviation of each gene’s weighted degree across all samples.

### 3.4 Predicting overall survival of lung adenocarcinoma patients using Weighted Gene Degrees

After having used unsupervised strategies to mine associations between SSN topology and lung adenocarcinoma heterogeneity, we set out to employ a supervised learning approach with the goal of predicting each patient’s overall survival using network based features. Specifically, we used WGDs as input for a Lasso regularized Cox proportional hazards regression model. In general, we benchmarked three different modeling scenarios: Model L, where LASSO was applied to the full set of WGDs; Model U+L, where LASSO was applied to a set of genes previously selected by a univariate filter; and Model U+L+R, which involved a relaxed fit of the same model.

To compare the global performance of the three above mentioned regression models, we computed both Harrel’s C-Index as well as the Integrated Time Dependent Brier’s Score for 100 models built using different training/testing data partitions (70% and 30% split, respectively). **Figure 7A** highlights the obtained C-Index values across the three frameworks. As expected, using a univariate cox regression as a filtering step prior to LASSO shrinkage significantly improved the predictive performance compared to the model using the full WGD matrix (Wilcoxon test, p-value *<* 0.05). On the other hand, models U+L and U+L+R show very similar predictive capabilities, with the unrelaxed fit showing a slightly higher median C-Index. Conversely, when assessing each model’s performance through the Integrated Brier’s Score statistic, no significant differences where found (**Figure 7B**). Overall, both metrics for evaluating survival predictions show that the three models perform better than at random in the majority of scenarios.

**Fig. 7.**
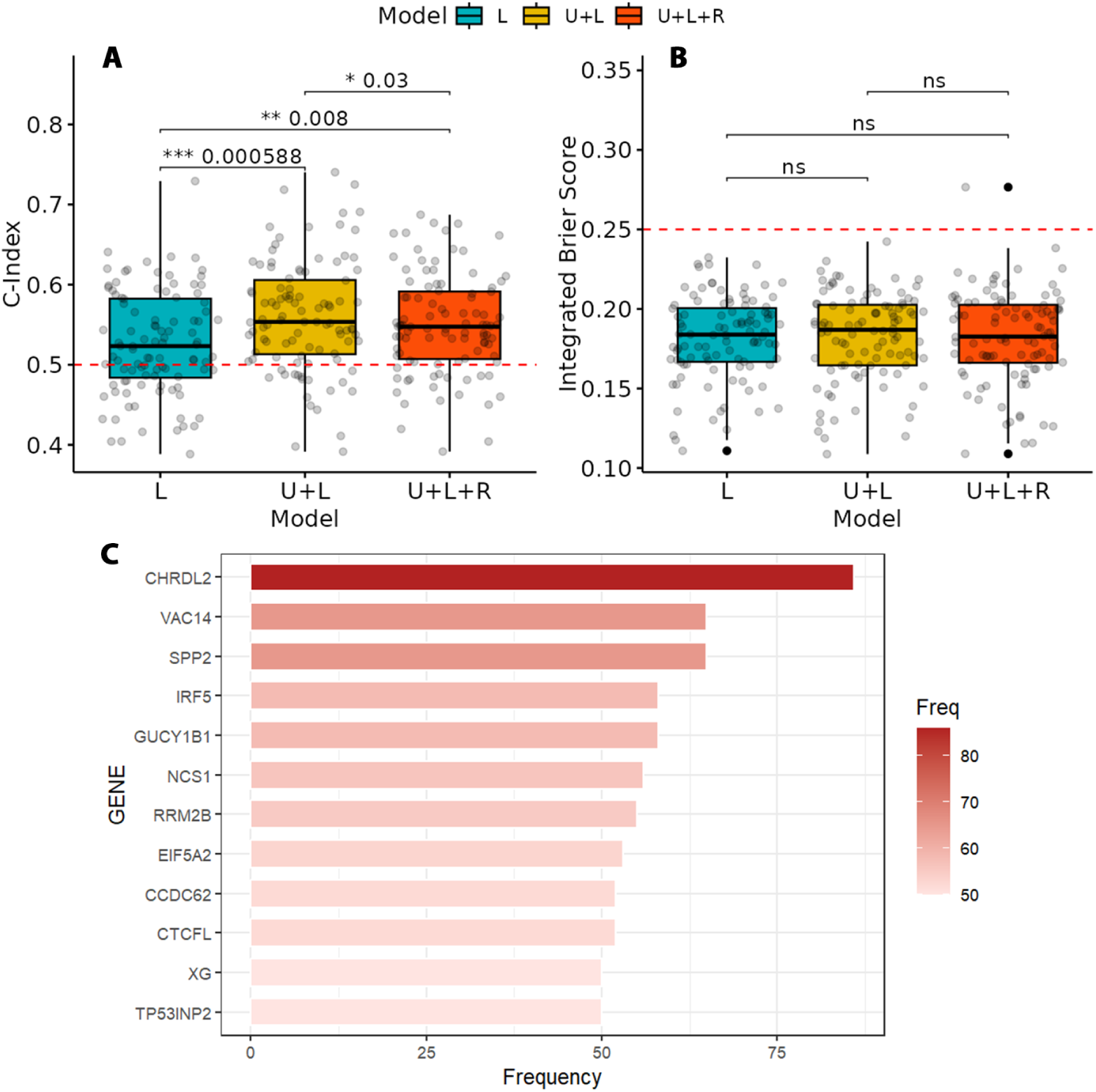
Overall predictive performance, uncertainty, and stability of Lasso gene selection in 100 random training/testing data partitions. 3 modelling frameworks were contrasted: Model L, where Lasso is applied to the entire set of genes, Model U+L, where a univariate filter is applied a priori to Lasso, and model U+L+R, where the fit obtained on the U+L model is relaxed (known as relaxed Lasso). **A-B:** Overall performance and uncertainty measured by **A**, Concordance Index (C-Index), and **B**, Integrated Brier Score. Red dotted lines mark the score in performance where model prediction are indistinct from a random guesser. Performance differences were calculated through Wilcoxon signed rank tests. **C:** Set of genes considered stable predictors of survival. Each gene is plotted against the frequency with which they were selected by both the univariate and Lasso filters in the random resampling procedure.

One of the main advantages of using regularized regression strategies is their ability to perform feature selection while reducing multicollinearity in high dimensional case scenarios such as those commonly found in “omics” data sets. However, given how LASSO selects a random representative variable among a group of correlated predictors [38], it is possible that dissimilar sets of genes are selected in different iterations of the model fitting process. Thus, we consider a gene’s weighted degree to be stably associated to survival only if it is selected by both the univariate filter and LASSO in at least 50 of the 100 random data partitions. Notably, a group of 12 genes were selected to be stably associated with patient survival, with Chordin Like 2 (CHRDL2) being the gene most frequently picked, followed by a tie between VAC14 Component of PIKFYVE Complex (VAC14) and Secreted Phosphoprotein 2 (SPP2) (**Figure 7C**). **Supplementary Figure 3** shows the association of all stable genes’ weighted degrees with overall patient survival.

Additionally, we focused on a representative U+L+R model to assess overall goodness of fit and local predictive performance on a discrete scale, as it was the best performing model among the three frameworks in a random iteration. The obtained model contained 65 genes, of which 7 were considered stable by the resampling procedure described above. Among the stable features, CHRDL2 and SPP2 are the two genes with the largest negative and positive coefficients, respectively (**Figure 8A**). To validate the sign of association between both genes weighted degree and survival, we stratified the full data set by their median weighted degree. In both cases, the groups with high and low weighted degrees correspond with the sign of the coefficients obtained in the model and show significant differences in overall survival probabilities (**Figure 8D-E**). When evaluating the full model, we observed a significant difference in the survival distribution of both risk groups obtained by stratifying the linear predictors of the unseen test data (**Figure 8B**). Moreover, the assessment of 1-, 2-, 3-, and 5-year overall survival prediction yielded AUC scores of 0.63, 0.63, 0.64, and 0.67, respectively (**Figure 8F**). Finally, the time-dependent Brier score on all possible time-points showed a progressive loss of predictive performance over time, reaching an asymptotic behavior after the 1000-day mark, likely due to the lack of events with longer time spans (**Figure 8C**).

**Fig. 8.**
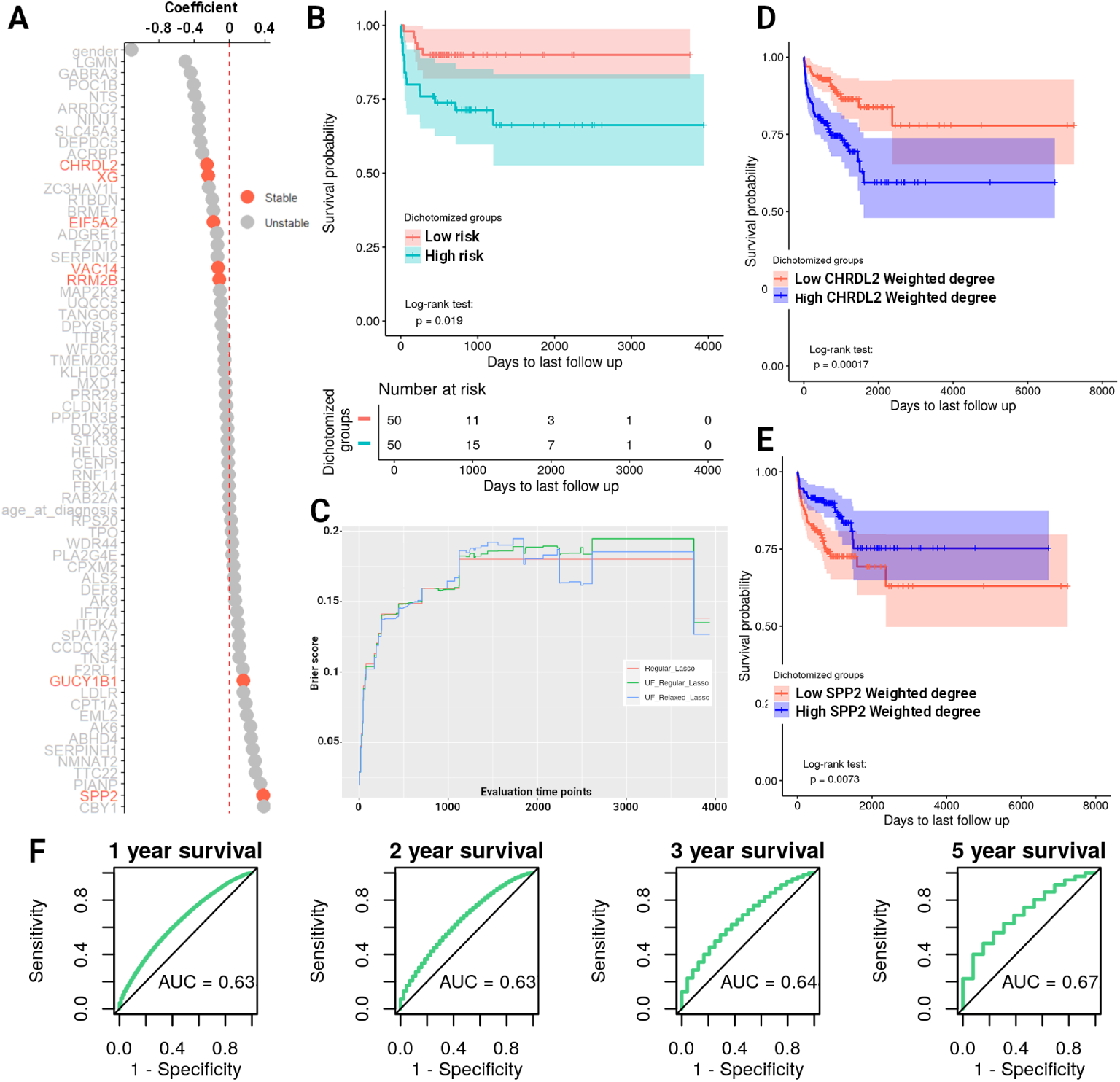
Model diagnostics of a representative relaxed model. **A:**Coefficient plot of 65 genes selected after univariate and Lasso filtering. Genes hilighted in orange were considered to be stable predictors. **B:**Kaplan-Meier plot of the stratified risk scores predicted for the test data set using the relaxed fit. **C:** Time-Dependent Brier Score curve for all available time points in the test data set. **D-E:**Kaplan-Meier plots of CHRDL2 and SPP2 stratified by their median weighted degree in the entire cohort. **F:** AUC scores for prediction of 1-,2-,3-,5-year overall survival in the test data.

## 4 Discussion

This work aimed to characterize the link between GCN topology and heterogeneous phenotypes of patients with lung adenocarcinoma. To this end, we modelled single sample GCNs for patients in the LUAD cohort of the TCGA data base and analyzed them through both unsupervised and supervised methods. Here we focused on patientspecific network similarity and node weighted degree centrality as the topological features used to identify patterns in the co-expression landscape at a population level.

When focusing on shared gene interactions, a clear community structure can be observed in the patient-similarity graph. As previously mentioned, differing clinical backgrounds were not sufficient to explain the observed grouping. Nevertheless, Cluster 1 was one of two communities with a significant overrepresentation of a clinical feature, being specifically enriched with T1 tumors (i.e., growths ≤ 3 cm surrounded by lung or pleura with no local invasion;[39]). Concomitantly, Cluster 1 had the consensus network with the highest degree of conservation among its patients and displayed enriched functional processes associated with cilium genesis, structure, and maintenance. Ciliated cells are considered the most predominant cell type in the lung airway and the most terminally differentiated, showing very limited proliferative potential [40]. Furthermore, ciliated cells have been proposed as one of the cell types capable of forming precursory lesions that trigger the origin of pulmonary LUAD tumors [41]. Together, this suggests that tumors sampled from patients in Cluster 1 are likely to represent a cancerous phenotype more indicative of the earlier stages of progression compared to the rest of the cohort, as the SSNs of these patients illustrate the regular transcriptional activity that is most conserved in ciliated lung epithelia.

On the other hand, networks in Cluster 5, which showed a significant overrepresentation of female patients, are primarily composed of genes involved in multiple biological processes related to cell cycle regulation and progression. Gender biases have been observed in lung cancer development, response to treatment, and mortality [42]. Specifically, the link between female sex hormones and cell proliferation programs in cancer has been previously reported. For example, not only is estrogen capable of inducing cell proliferation in lung cancer, but it can also be actively produced by NSCLC cells [43]. Conversely, progesterone has been shown to inhibit lung cancer growth in mice, while Progesterone Receptor-positive (PR+) NSCLCs in female patients frequently present an inverse correlation between PR activity and overall disease progression [44]. Additionally, sex-biased genetic factors are well established to play a role in the progression of lung cancer. For instance, female patients are more likely to have driver mutations in genes such as EGFR or KRAS [45]. Similarly, a meta-analysis of gender-biased somatic mutations in the TCGA LUAD cohort found that, while EGFR did not meet the significance threshold established by the authors, it displayed a higher mutation frequency in female patients consistent with the literature. The same study found that MED12 was one of 5 genes with a female-biased mutation signature [46]. Interestingly, EGFR and MED12 are exclusive components of Cluster’s 5 consensus network, despite MED12 not being annotated in a biological process associated with cell proliferation. This suggests that gender-biased mutation patterns could also be contributing to the overall structure of SSN’s architecture. In summary, our analysis highlights a conserved gene co-expression signature associated to tumor growth and proliferation, predominantly among female patients, that likely reflects the cumulative impact that heterogenous factors contribute to gender differences in LUAD patient outcomes.

Next, owing to the pattern of shared of biological programs associated to consensus networks of Cluster 3 and Cluster 6, we identified significant differences in the overall tumor microenvironment composition across all patient communities. Above all, Cluster 3 consistently showed a depletion of both immune and stromal cell infiltrates, whereas Cluster 6 displayed the opposite pattern, with frequent high levels of immune infiltration. Additionally, Clusters 1 and 5 displayed the highest levels of endothelial cell composition, consistent with our previously stated hypothesis of Cluster 1 being primarily composed of earlier-staged tumors, as it is known that lung endothelial cells comprise the pulmonary vascular bed that is needed for gas exchange to take place [47]. Furthermore, while Cluster 2 portrayed a unique signature more associated with cell types that exert pro-tumoral effects, such as Th2 cells [48] and M2 macrophages [49], Cluster 6 showed a preferential enrichment of cells that elicit anti-tumoral activity, such as Macrophages M1[49], B-cells [50][51], and CD8+ Tcm and Tem cells[52].

Notwithstanding, our analysis was not able to identify any significant association between the groups with high and low immune infiltration and patient clinical outcomes. This is likely due to the fact that the relationship between immune infiltration and patient prognosis varies in a context specific manner, thus reflecting the functional plasticity displayed by immune system cells in the tumor microenvironment. However, the establishment immunosuppressive phenotypes are known to be a crucial event that favours tumor malignancy. Hence, the identification of clusters of patients with diverse immune system activity highlights the potential of SSNs as a tool for probing the network of genes that drive immunosuppressive states, thereby warranting further research.

Turning to another aspect of our study, we investigated whether the weighted degrees of genes in SSNs could be a topological feature of utility for downstream analyses. Of note, previous studies have tried a similar approach wherein the degree of nodes in correlation-based SSNs (i.e. co-expression networks) is used as a summary statistic to uncover biologically relevant associations. For example, Chen et. al. performed SSN modelling using the SWEET method (a SSN inference method based on Pearson Correlation and genome-wide sample weights) and built a Network Degree Matrix to study whether SSN degree distributions were correlated with genetic dependency scores of 580 cell lines. More specifically in the LUAD-TCGA cohort, they utilized the same approach to discover “dark” genes with degree differences between two defined subtypes that were not explained by differential expression analysis [53]. Another example of a similar network-analytical approach was employed by Dai et al., whereby cell-specific networks built using a mutual dependency criterion to establish gene-gene correlations were further summarized by calculating a Network Degree Matrix for downstream analyses, analogous to standard single-cell RNA-Seq pipelines [54].

To our knowledge, this work is the first to employ the same network analysis strategy using LIONESS-based fully-connected mutual information networks. As a straightforward proof of concept of the feasibility of our constructed Degree Matrix, we performed GSEA on each gene degree’s standard deviation to assess whether this network metric was indeed reflective of biologically relevant signals. Our results indicated that degree variation within LUAD SSNs is significantly associated to cell cycle regulatory programs such as the hallmarks Myc Targets V1, Myc targets V2, and E2F targets as well as multiple biological processes consistent with the overall trend (see **Figure 6**). Alterations in the MYC proto-oncogene regulatory network have been observed to be present in nearly 40% of human cancers, while whole-exome sequencing studies have found its amplifications to be present in approximately one third of LUADs [55]. Similarly, the E2F family of transcription factors are well established to be involved in several types of cancer [56]. Specifically in LUAD, independent studies have found the expression of E2F2 and E2F7 to be prognostic for patient survival and disease progression by promoting cell proliferative programs [57] [56]. Therefore, we can assume that degrees obtained from Mutual Information based SSNs are capable of depicting known aspects of lung cancer biology, and could potentially benefit the search for novel biomarkers of patient prognosis.

Next, working under the driving hypothesis that SSN weighted degrees are enriched with biological information, we sought to investigate whether they could further be predictive of LUAD patient survival. To this end, we used gene weighted degrees as input for regularized Cox proportional hazards regression approaches and validated the model building procedure and performance through random resampling methods. The proposed methodology highlighted 12 genes whose structural importance in individual GCNs is robustly associated to survival, regardless of the established training and testing data partition. Notably, all 12 genes (CHRDL2, SPP2, VAC14, IRF5, GUCY1B1, NCS1, RRM2B, EIF5A2, CCDC62, CTCFL, XG and TP53INP2) have been previously associated to various types of cancer [58] [59] [60] [61] [62] [63] [64] [65] [66] [67] [68] [69], with 7 (CHRDL2, SPP2, IRF5, NCS1, RRM2B, EIF5A2, CTCFL) including reports specific to lung cancer [70] [59] [71] [72] [73] [74] [75].

It is worth mentioning that the functional assessment of sets of genes selected by LASSO based on gene enrichment analysis may be hindered by the removal of collinearity between variables in the model fitting step. However, there is an interesting overlap in the body of literature that places these genes in the context of cancer research. For example, CHRDL2 and SPP2, the two most stable predictors of survival in this analysis, have been shown to mainly operate as antagonists of Bone Morphogenetic Proteins (BMPs), a group of growth factor cytokines that belong to the transforming growth factor *β* (TGF-*β*) superfamily of proteins. While the baseline interactions between CHRDL2/SPP2 and the BMP signalling pathway have been more extensively described in the context of bone development and homeostasis, multiple studies have linked both genes to cancer-promoting processes mediated by BMP signalling. For example, overexpression of CHRDL2 has been shown to promote proliferation and metastasis in osteosarcoma tissues and cell lines by blocking the BMP-9/PI3K/AKT pathway [76]. Similarly, in colorectal cancer, CHRDL2 overexpression promotes cell proliferation and inhibits apoptosis by blocking BMP2-driven Smad1/5 signaling [77]. Conversely, BMP2 blockade via SPP2 administration has been shown to inhibit tumor cell growth in hepatocellular carcinoma [78], osteosarcoma [79], prostate cancer [80], pancreatic cancer [81] and lung cancer [82].

The role of BMP signaling in NSCLC growth [83], and more recently in LUAD metastasis and cell differentiation [84] has been demonstrated to be of great relevance. However, added layers of complexity are known to take part in the BMP-induced carcinogenic process. One such case is the crosstalk between BMP, TGF-*β* and WNT/*β*-catenin pathways, which together contribute to the formation of cancerrelated phenotypes [85]. We hypothesize that other genes highlighted in our analysis, such as EIF5A, TP53INP2 and CTCFL/BORIS are reflective of this crossroad between signalling routes. For instance, EIF5A has been shown to promote epithelial to mesenchymal transition (EMT), treatment resistance and metastasis through direct dysregulation of TGF-*β* signaling in a variety of cancers [86] [87] [88], [89]. TP53INP2, an autophagy-related protein, was observed to modulate EMT, migration and invasion of bladder cancer cells [90], as well as the malignant progression of colorectal cancer [69] through *β*-catenin dependent signalling . In parallel, CTCFL/BORIS, a crucial regulator of the malignant effects of cancer stem cells, is known to primarily exert its tumorigenic actions through both Wnt/*β*-catenin and NOTCH pathways [91]. However, TGF-*β* dependent signalling of CTCFL/BORIS has also been observed to be relevant in melanoma invasiveness [92], and neuroblastoma migration [93].

Together, given that SSN analysis is capable of representing traits of the population’s underlying heterogeneity as structural features of transcriptome-wide GCNs, we believe that the underscored set of genes associated to survival indeed portray the complex and context-specific nature of the transcriptional regulation that BMP, TGF-*β* and WNT/*β*-catenin signaling pathways may exert to influence LUAD clinical outcomes. However, further research needs to be addressed to support this hypothesis.

## 5 Conclusions

Cancer is a putative heterogenous disease. A myriad of studies addressing the discrete relationship between clinical-, cellular- and molecular-context specificity have motivated the development of new analytical strategies capable of embracing such phenotypic complexity. To this end, the use of patient-specific GCNs could aid bridge the gap between the studies of minute changes in transcriptional activity/regulation and cancer heterogeneity.

In this work, we applied unsupervised and supervised methods to analyze changes in the co-expression patterns at an individual level for patients with LUAD. We hereby summarise the two principal findings of this study:

1. Using a metric of SSN similarity between patients, a graph-clustering strategy highlighted 6 different subtypes of LUAD. We found that clinical phenotypes were not the main driver of SSN similarity, however, two clusters (Cluster 1 and Cluster 5) displayed a significant enrichment of specific tumor classification and gender. On the other hand, cellular and molecular phenotypes were more clearly divisive between clusters, showing very distinct immune infiltration composition as well as enriched biological processes in their most conserved edges.
2. By calculating the weighted degree of each gene in an individual network, we observed that relevant biological signals consistent with known cancer biology can be retrieved. Furthermore, by using this node-centrality metric, we were able to capture complex traits related to each gene’s structural role in a network and use them as features in a supervised learning scheme with the goal of finding novel network-based predictors of survival. Our approach underscored 12 genes (CHRDL2, SPP2, VAC14, IRF5, GUCY1B1, NCS1, RRM2B, EIF5A2, CCDC62, CTCFL, XG and TP53INP2) with known associations to the origin and progression of cancer, whose topological relevance in a network is predictive of patient survival in LUAD.

Regarding the limitations of our research, it should be stressed that all of these analyses were performed using the TCGA-LUAD cohort only. Given that SSN’s place population heterogeneity as the main subject of study, new workflows integrating multiple sources of variation in every step of the procedure, including the individual network building stage, should be considered in order to validate that the results here obtained are not strictly dependent on the choice of data set. However, such a strategy involving multiple data sets and larger sample sizes are posed to be an expensive computational task when using mutual information as a network building method. Thus, careful experimental design should be taken into account in regards to the specific biological questions at hand.

The choice of an “ideal” set of background samples from which SSNs will be based on, is a major challenge of multiple SSN building methods that has been previously raised [94]. While the resampling methods employed in this work are effective means to attenuate data-dependent effects in predictive modelling workflows, an adequate independent validation analysis of the employed unsupervised and supervised methods needs to be pursued. Thus, we envision expanding our analysis pipeline to the entire NSCLC landscape as well as cancers with similar clinical outcomes as a first perspective avenue.

Finally, we contend that not only will the workflow implemented in this study significantly contribute to the growing methodological and analytical toolkit that patient-specific network understanding demands, but also that the biological insights uncovered will guide future endeavors in the fields of network-medicine and lung cancer research.

## Supplementary information

**Table 1.** Structural summary statistics of per-cluster consensus networks.

| Global Descriptor | Cluster 1 | Cluster 2 | Cluster 3 | Cluster 4 | Cluster 5 | Cluster 6 |
| --- | --- | --- | --- | --- | --- | --- |
| Number of nodes | 400 | 491 | 774 | 2257 | 528 | 1114 |
| Number of edges | 1394 | 1381 | 2360 | 4767 | 2599 | 2108 |
| Avg. Number of neighbors | 6.9 | 5.8 | 6.2 | 6.2 | 10.1 | 3.8 |
| Network diameter | 7 | 9 | 9 | 15 | 6 | 12 |
| Characteristic path length | 3.4 | 3.6 | 3.5 | 4.6 | 3 | 4.5 |
| Clustering coefficient | 0.056 | 0.084 | 0.087 | 0.089 | 0.102 | 0.039 |
| Network density | 0.017 | 0.013 | 0.008 | 0.005 | 0.02 | 0.004 |
| Network centralization | 0.106 | 0.153 | 0.101 | 0.081 | 0.145 | 0.048 |
| Number of connected components | 1 | 11 | 9 | 347 | 7 | 21 |

**Fig. 1.**
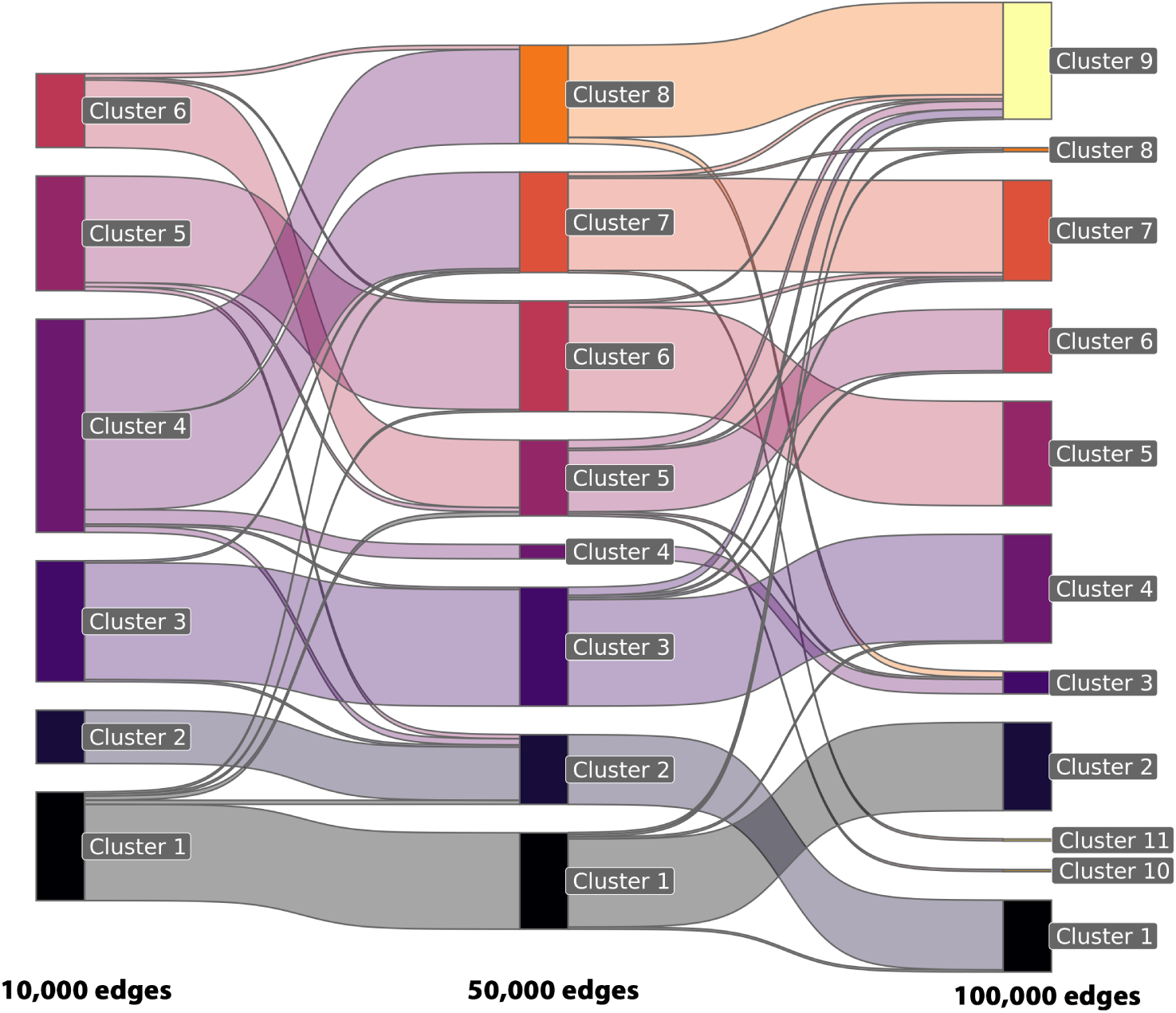
Sankey diagram illustrating the agreement between clustering results based on patient-specific network similarity at increasing network sizes. Louvain clustering for weighted graphs was used in all cases.

**Fig. 2.**
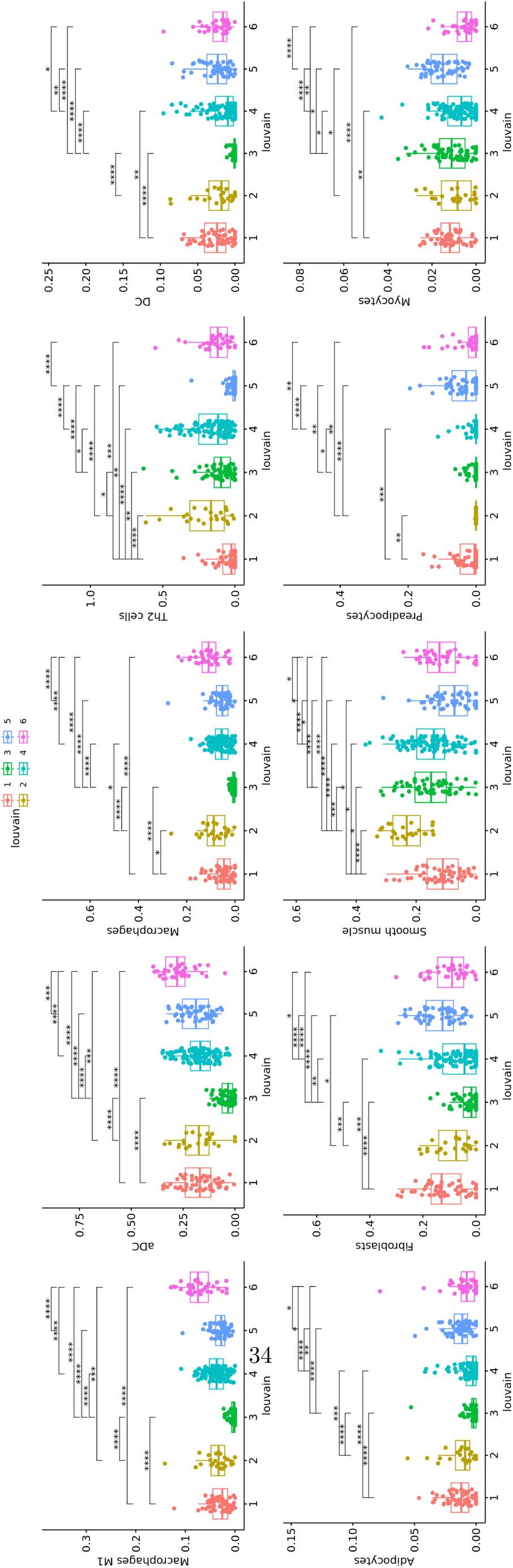
Box-plots showing the distribution of the Top-5 Xcell Immune and Stromal cell types, according to the significance obtained in the Kruskal-Wallis test osberved in Figure 5A

**Fig. 3.**
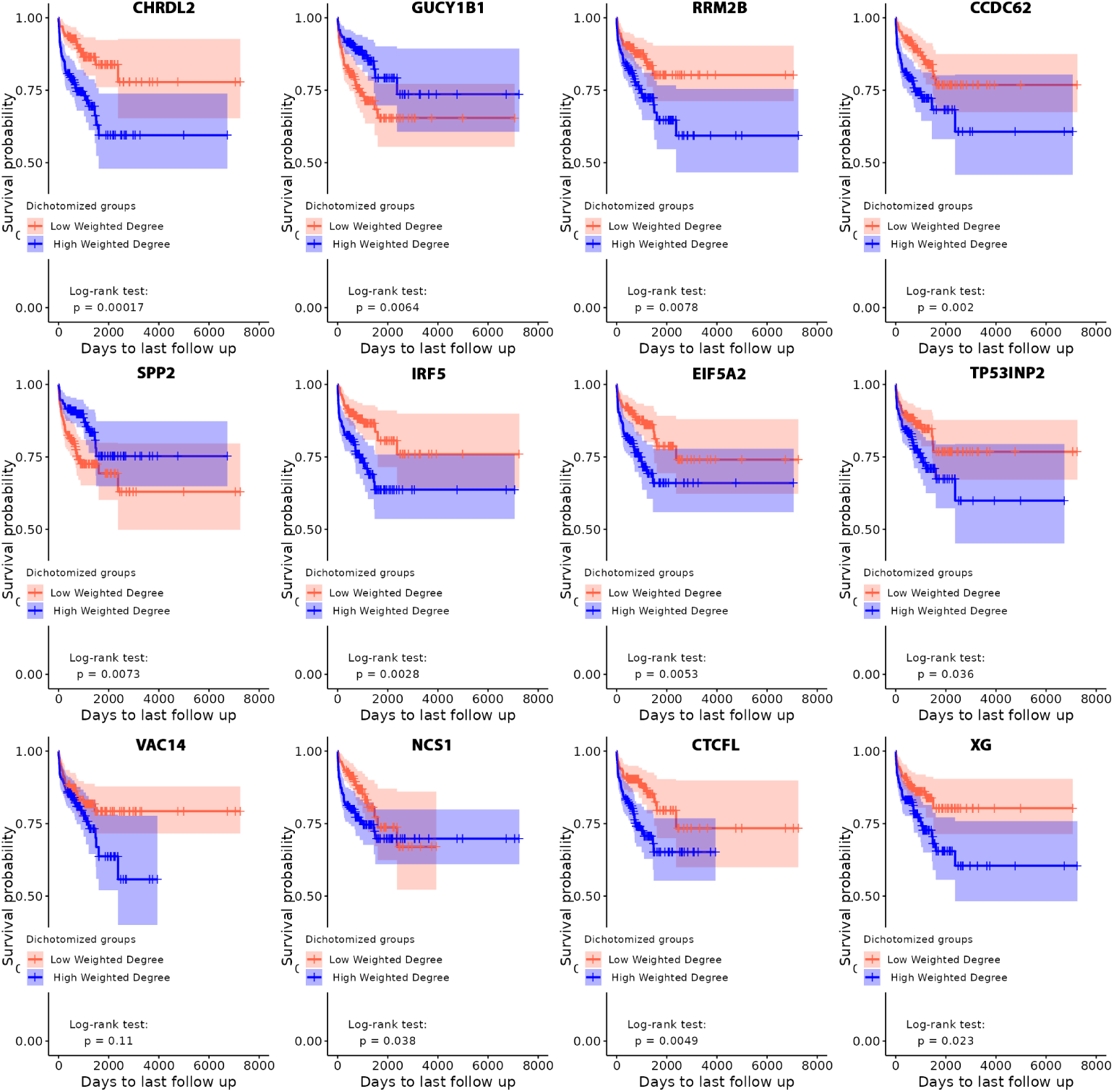
Kaplan-Meier plots of the 12 stable predictors highlighted in our analysis stratified by their median weighted degrees in the entire cohort.

## References

[1] Sung, H., Ferlay, J., Siegel, R.L., Laversanne, M., Soerjomataram, I., Jemal, A., Bray, F.: Global Cancer Statistics 2020: GLOBOCAN Estimates of Incidence and Mortality Worldwide for 36 Cancers in 185 Countries. CA: a cancer journal for clinicians 71(3), 209–249 (2021) 10.3322/caac.21660

[2] Perez-Moreno, P., Brambilla, E., Thomas, R., Soria, J.-C.: Squamous Cell Carcinoma of the Lung: Molecular Subtypes and Therapeutic Opportunities. Clinical Cancer Research 18(9), 2443–2451 (2012) 10.1158/1078-0432.CCR-11-2370

[3] Wang, B.-Y., Huang, J.-Y., Chen, H.-C., Lin, C.-H., Lin, S.-H., Hung, W.-H., Cheng, Y.-F.: The comparison between adenocarcinoma and squamous cell carcinoma in lung cancer patients. Journal of Cancer Research and Clinical Oncology 146(1), 43–52 (2020) 10.1007/s00432-019-03079-8

[4] Bade, B.C., Dela Cruz, C.S.: Lung Cancer 2020: Epidemiology, Etiology, and Prevention. Clinics in Chest Medicine 41(1), 1–24 (2020) 10.1016/j. ccm.2019.10.001

[5] Du, W., Elemento, O.: Cancer systems biology: Embracing complexity to develop better anticancer therapeutic strategies. Oncogene 34(25), 3215–3225 (2015) 10.1038/onc.2014.291

[6] Serin, E.A.R., Nijveen, H., Hilhorst, H.W.M., Ligterink, W.: Learning from Coexpression Networks: Possibilities and Challenges. Frontiers in Plant Science 7 (2016) 10.3389/fpls.2016.00444

[7] Espinal-Enríquez, J., Fresno, C., Anda-Jáuregui, G., Hernández-Lemus, E.: RNA-Seq based genome-wide analysis reveals loss of inter-chromosomal regulation in breast cancer. Scientific Reports 7(1), 1760 (2017) 10.1038/s41598-017-01314-1

[8] García-Cortés, D., de Anda-Jáuregui, G., Fresno, C., Hernández-Lemus, E., Espinal-Enríquez, J.: Gene Co-expression Is Distance-Dependent in Breast Cancer. Frontiers in Oncology 10 (2020) 10.3389/fonc.2020.01232

[9] Zamora-Fuentes, J.M., Hernández-Lemus, E., Espinal-Enríquez, J.: Gene Expression and Co-expression Networks Are Strongly Altered Through Stages in Clear Cell Renal Carcinoma. Frontiers in Genetics 11, 578679 (2020) 10.3389/fgene.2020.578679

[10] Andonegui-Elguera, S.D., Zamora-Fuentes, J.M., Espinal-Enríquez, J., Hernández-Lemus, E.: Loss of Long Distance Co-Expression in Lung Cancer. Frontiers in Genetics 12 (2021) 10.3389/fgene.2021.625741

[11] Nakamura-García, A.K., Espinal-Enríquez, J.: The network structure of hematopoietic cancers. Scientific Reports 13(1), 19837 (2023) 10.1038/s41598-023-46655-2

[12] Kuijjer, M.L., Tung, M.G., Yuan, G., Quackenbush, J., Glass, K.: Estimating Sample-Specific Regulatory Networks. iScience 14, 226–240 (2019) 10.1016/j.isci.2019.03.021

[13] Colaprico, A., Silva, T.C., Olsen, C., Garofano, L., Cava, C., Garolini, D., Sabedot, T.S., Malta, T.M., Pagnotta, S.M., Castiglioni, I., Ceccarelli, M., Bontempi, G., Noushmehr, H.: TCGAbiolinks: An R/Bioconductor package for integrative analysis of TCGA data. Nucleic Acids Research 44(8), 71 (2016) 10.1093/nar/gkv1507

[14] Margolin, A.A., Nemenman, I., Basso, K., Wiggins, C., Stolovitzky, G., Favera, R.D., Califano, A.: ARACNE: An Algorithm for the Reconstruction of Gene Regulatory Networks in a Mammalian Cellular Context. BMC Bioinformatics 7(1), 7 (2006) 10.1186/1471-2105-7-S1-S7

[15] Csardi, G., Nepusz, T.: The igraph software package for complex network research. InterJournal **Complex Systems**, 1695 (2006)

[16] Csárdi, G., Nepusz, T., Traag, V., Horvát, S., Zanini, F., Noom, D., Müller, K.: igraph: Network Analysis and Visualization in R. (2024). 10.5281/zenodo.7682609. R package version 2.0.3. https://CRAN.R-project.org/package=igraph

[17] Lancichinetti, A., Fortunato, S.: Limits of modularity maximization in community detection. Physical Review E 84(6), 066122 (2011) 10.1103/PhysRevE.84.066122 arXiv:1107.1155 [physics]

[18] Aran, D.: Cell-Type Enrichment Analysis of Bulk Transcriptomes Using xCell. Methods in Molecular Biology (Clifton, N.J.) 2120, 263–276 (2020) 10.1007/978-1-0716-0327-71_9

[19] Morselli Gysi, D., de Miranda Fragoso, T., Zebardast, F., Bertoli, W., Busskamp, V., Almaas, E., Nowick, K.: Whole transcriptomic network analysis using Coexpression Differential Network Analysis (CoDiNA). PloS One 15(10), 0240523 (2020) 10.1371/journal.pone.0240523

[20] Morris, J.H., Apeltsin, L., Newman, A.M., Baumbach, J., Wittkop, T., Su, G., Bader, G.D., Ferrin, T.E.: clusterMaker: A multi-algorithm clustering plugin for Cytoscape. BMC Bioinformatics 12(1), 436 (2011) 10.1186/1471-2105-12-436

[21] Kolberg, L., Raudvere, U., Kuzmin, I., Adler, P., Vilo, J., Peterson, H.: G:Profiler—interoperable web service for functional enrichment analysis and gene identifier mapping (2023 update). Nucleic Acids Research 51(W1), 207–212 (2023) 10.1093/nar/gkad347

[22] Shannon, P., Markiel, A., Ozier, O., Baliga, N.S., Wang, J.T., Ramage, D., Amin, N., Schwikowski, B., Ideker, T.: Cytoscape: A Software Environment for Integrated Models of Biomolecular Interaction Networks. Genome Research 13(11), 2498-2504 (2003) 10.1101/gr.1239303

[23] Lopes-Ramos, C.M., Belova, T., Brunner, T.H., Ben Guebila, M., Osorio, D., Quackenbush, J., Kuijjer, M.L.: Regulatory Network of PD1 Signaling Is Associated with Prognosis in Glioblastoma Multiforme. Cancer Research 81(21), 5401–5412 (2021) 10.1158/0008-5472.CAN-21-0730

[24] Belova, T., Biondi, N., Hsieh, P.-H., Lutsik, P., Chudasama, P., Kuijjer, M.L.: Heterogeneity in the gene regulatory landscape of leiomyosarcoma. NAR Cancer 5(3), 037 (2023) 10.1093/narcan/zcad037

[25] Lopes-Ramos, C.M., Kuijjer, M.L., Ogino, S., Fuchs, C.S., DeMeo, D.L., Glass, K., Quackenbush, J.: Gene Regulatory Network Analysis Identifies Sex-Linked Differences in Colon Cancer Drug Metabolism. Cancer Research 78(19), 5538–5547 (2018) 10.1158/0008-5472.CAN-18-0454

[26] Subramanian, A., Tamayo, P., Mootha, V.K., Mukherjee, S., Ebert, B.L., Gillette, M.A., Paulovich, A., Pomeroy, S.L., Golub, T.R., Lander, E.S., Mesirov, J.P.: Gene set enrichment analysis: A knowledge-based approach for interpreting genome-wide expression profiles. Proceedings of the National Academy of Sciences of the United States of America 102(43), 15545–15550 (2005) 10.1073/pnas.0506580102

[27] Mootha, V.K., Lindgren, C.M., Eriksson, K.-F., Subramanian, A., Sihag, S., Lehar, J., Puigserver, P., Carlsson, E., Ridderstråle, M., Laurila, E., Houstis, N., Daly, M.J., Patterson, N., Mesirov, J.P., Golub, T.R., Tamayo, P., Spiegelman, B., Lander, E.S., Hirschhorn, J.N., Altshuler, D., Groop, L.C.: PGC-1*α*-responsive genes involved in oxidative phosphorylation are coordinately downregulated in human diabetes. Nature Genetics 34(3), 267–273 (2003) 10.1038/ ng1180

[28] Ashburner, M., Ball, C.A., Blake, J.A., Botstein, D., Butler, H., Cherry, J.M., Davis, A.P., Dolinski, K., Dwight, S.S., Eppig, J.T., Harris, M.A., Hill, D.P., Issel-Tarver, L., Kasarskis, A., Lewis, S., Matese, J.C., Richardson, J.E., Ringwald, M., Rubin, G.M., Sherlock, G.: Gene Ontology: Tool for the unification of biology. Nature Genetics 25(1), 25–29 (2000) 10.1038/75556

[29] Liberzon, A., Birger, C., Thorvaldsdóttir, H., Ghandi, M., Mesirov, J.P., Tamayo, P.: The Molecular Signatures Database (MSigDB) hallmark gene set collection. Cell Systems 1(6), 417–425 (2015) 10.1016/j.cels.2015.12.004

[30] Tibshirani, R.: The lasso method for variable selection in the Cox model. Statistics in Medicine 16(4), 385–395 (1997) 10.1002/(sici)1097-0258(19970228)16:4⟨385::aid-sim380⟩3.0.co;2-3

[31] Tibshirani, R.: Regression Shrinkage and Selection via the Lasso. Journal of the Royal Statistical Society. Series B (Methodological) 58(1), 267–288 (1996) 2346178

[32] Ying, X.: An Overview of Overfitting and its Solutions. Journal of Physics: Conference Series 1168, 022022 (2019) 10.1088/1742-6596/1168/2/022022

[33] Su, W., Bogdan, M., Candes, E.: False Discoveries Occur Early on the Lasso Path. arXiv (2016). 10.48550/arXiv.1511.01957

[34] Meinshausen, N.: Relaxed Lasso. Computational Statistics & Data Analysis 52(1), 374–393 (2007) 10.1016/j.csda.2006.12.019

[35] Friedman, J.H., Hastie, T., Tibshirani, R.: Regularization Paths for Generalized Linear Models via Coordinate Descent. Journal of Statistical Software 33, 1–22 (2010) 10.18637/jss.v033.i01

[36] Simon, N., Friedman, J., Hastie, T., Tibshirani, R.: Regularization Paths for Cox’s Proportional Hazards Model via Coordinate Descent. Journal of Statistical Software 39(5), 1–13 (2011) 10.18637/jss.v039.i05

[37] Harrell, F.E.: Evaluating the Yield of Medical Tests. JAMA: The Journal of the American Medical Association 247(18), 2543 (1982) 10.1001/jama.1982.03320430047030

[38] Freijeiro-González, L., Febrero-Bande, M., González-Manteiga, W.: A Critical Review of LASSO and Its Derivatives for Variable Selection Under Dependence Among Covariates. International Statistical Review 90(1), 118–145 (2022) 10.1111/insr.12469

[39] Lababede, O., Meziane, M.A.: The Eighth Edition of TNM Staging of Lung Cancer: Reference Chart and Diagrams. The Oncologist 23(7), 844–848 (2018) 10.1634/theoncologist.2017-0659

[40] Sainz de Aja, J., Dost, A.F.M., Kim, C.F.: Alveolar progenitor cells and the origin of lung cancer. Journal of Internal Medicine 289(5), 629–635 (2021) 10.1111/joim.13201

[41] Park, W.Y., Kim, M.H., Shin, D.H., Lee, J.H., Choi, K.U., Kim, J.Y., Park, D.Y., Lee, C.H., Sol, M.Y.: Ciliated adenocarcinomas of the lung: A tumor of non-terminal respiratory unit origin. Modern Pathology 25(9), 1265–1274 (2012) 10.1038/modpathol.2012.76

[42] May, L., Shows, K., Nana-Sinkam, P., Li, H., Landry, J.W.: Sex Differences in Lung Cancer. Cancers 15(12), 3111 (2023) 10.3390/cancers15123111

[43] Pietras, R.J., Márquez, D.C., Chen, H.-W., Tsai, E., Weinberg, O., Fishbein, M.: Estrogen and growth factor receptor interactions in human breast and non-small cell lung cancer cells. Steroids 70(5-7), 372–381 (2005) 10.1016/j.steroids.2005.02.017

[44] Ishibashi, H., Suzuki, T., Suzuki, S., Niikawa, H., Lu, L., Miki, Y., Moriya, T., Hayashi, S.-i., Handa, M., Kondo, T., Sasano, H.: Progesterone Receptor in Non– Small Cell Lung Cancer—A Potent Prognostic Factor and Possible Target for Endocrine Therapy. Cancer Research 65(14), 6450–6458 (2005) 10.1158/0008-5472.CAN-04-3087

[45] Florez, N., Kiel, L., Riano, I., Patel, S., DeCarli, K., Dhawan, N., Franco, I., Odai-Afotey, A., Meza, K., Swami, N., Patel, J., Sequist, L.V.: Lung Cancer in Women: The Past, Present, and Future. Clinical Lung Cancer 25(1), 1–8 (2024) 10.1016/j.cllc.2023.10.007

[46] Yuan, Y., Liu, L., Chen, H., Wang, Y., Xu, Y., Mao, H., Li, J., Mills, G.B., Shu, Y., Li, L., Liang, H.: Comprehensive Characterization of Molecular Differences in Cancer between Male and Female Patients. Cancer Cell 29(5), 711–722 (2016) 10.1016/j.ccell.2016.04.001

[47] Tsikis, S.T., Hirsch, T.I., Fligor, S.C., Quigley, M., Puder, M.: Targeting the lung endothelial niche to promote angiogenesis and regeneration: A review of applications. Frontiers in Molecular Biosciences 9 (2022) 10.3389/ fmolb.2022.1093369

[48] Frafjord, A., Buer, L., Hammarström, C., Aamodt, H., Woldbæk, P.R., Brustugun, O.T., Helland, Å, Øynebråten, I., Corthay, A.: The Immune Landscape of Human Primary Lung Tumors Is Th2 Skewed. Frontiers in Immunology 12, 764596 (2021) 10.3389/fimmu.2021.764596

[49] Zhang, B., Yao, G., Zhang, Y., Gao, J., Yang, B., Rao, Z., Gao, J.: M2-Polarized tumor-associated macrophages are associated with poor prognoses resulting from accelerated lymphangiogenesis in lung adenocarcinoma. Clinics 66(11), 1879–1886 (2011) 10.1590/S1807-59322011001100006

[50] Schalper, K.A., Brown, J., Carvajal-Hausdorf, D., McLaughlin, J., Velcheti, V., Syrigos, K.N., Herbst, R.S., Rimm, D.L.: Objective Measurement and Clinical Significance of TILs in Non–Small Cell Lung Cancer. JNCI: Journal of the National Cancer Institute 107(3), 435 (2015) 10.1093/jnci/ dju435

[51] Kinoshita, T., Muramatsu, R., Fujita, T., Nagumo, H., Sakurai, T., Noji, S., Takahata, E., Yaguchi, T., Tsukamoto, N., Kudo-Saito, C., Hayashi, Y., Kamiyama, I., Ohtsuka, T., Asamura, H., Kawakami, Y.: Prognostic value of tumor-infiltrating lymphocytes differs depending on histological type and smoking habit in completely resected non-small-cell lung cancer. Annals of Oncology 27(11), 2117–2123 (2016) 10.1093/annonc/mdw319

[52] [52] Manjarrez-Orduño, N., Menard, L.C., Kansal, S., Fischer, P., Kakrecha, B., Jiang, C., Cunningham, M., Greenawalt, D., Patel, V., Yang, M., Golhar, R., Carman, J.A., Lezhnin, S., Dai, H., Kayne, P.S., Suchard, S.J., Bernstein, S.H., Nadler, S.G.: Circulating T Cell Subpopulations Correlate With Immune Responses at the Tumor Site and Clinical Response to PD1 Inhibition in Non-Small Cell Lung Cancer. Frontiers in Immunology 9 (2018) 10.3389/fimmu.2018. 01613

[53] Chen, H.-H., Hsueh, C.-W., Lee, C.-H., Hao, T.-Y., Tu, T.-Y., Chang, L.-Y., Lee, J.-C., Lin, C.-Y.: SWEET: A single-sample network inference method for deciphering individual features in disease. Briefings in Bioinformatics 24(2), 032 (2023) 10.1093/bib/bbad032

[54] Dai, H., Li, L., Zeng, T., Chen, L.: Cell-specific network constructed by singlecell RNA sequencing data. Nucleic Acids Research 47(11), 62 (2019) 10.1093/nar/gkz172

[55] Eftekhari Kenzerki, M., Ahmadi, M., Mousavi, P., Ghafouri-Fard, S.: MYC and non-small cell lung cancer: A comprehensive review. Human Gene 37, 201185 (2023) 10.1016/j.humgen.2023.201185

[56] Wang, Q., Liu, J., Cheang, I., Li, J., Chen, T., Li, Y., Yu, B.: Comprehensive Analysis of the E2F Transcription Factor Family in Human Lung Adenocarcinoma. International Journal of General Medicine 15, 5973–5984 (2022) 10.2147/IJGM.S369582

[57] Du, K., Sun, S., Jiang, T., Liu, T., Zuo, X., Xia, X., Liu, X., Wang, Y., Bu, Y.: E2F2 promotes lung adenocarcinoma progression through B-Myband FOXM1-facilitated core transcription regulatory circuitry. International Journal of Biological Sciences 18(10), 4151–4170 (2022) 10.7150/ijbs.72386

[58] Wang, L., Xu, W., Mei, Y., Wang, X., Liu, W., Zhu, Z., Ni, Z.: CHRDL2 promotes cell proliferation by activating the YAP/TAZ signaling pathway in gastric cancer. Free Radical Biology and Medicine 193, 158–170 (2022) 10.1016/j.freeradbiomed.2022.09.006

[59] Tu, Y., Chen, C., Fan, G.: Association between the expression of secreted phosphoprotein - related genes and prognosis of human cancer. BMC Cancer 19, 1230 (2019) 10.1186/s12885-019-6441-3

[60] Wang, C.-H., Wang, L.-K., Wu, C.-C., Chen, M.-L., Kuo, C.-Y., Shyu, R.-Y., Tsai, F.-M.: TIG1 Inhibits the mTOR Signaling Pathway in Malignant Melanoma Through the VAC14 Protein. Anticancer Research 43(6), 2635–2643 (2023) 10.21873/anticanres.16430

[61] Roberts, B.K., Collado, G., Barnes, B.J.: Role of interferon regulatory factor 5 (IRF5) in tumor progression: Prognostic and therapeutic potential. Biochimica et Biophysica Acta (BBA) - Reviews on Cancer 1879(1), 189061 (2024) 10.1016/j.bbcan.2023.189061

[62] Sadeghi, M., Karimi, M.R., Karimi, A.H., Ghorbanpour Farshbaf, N., Barzegar, A., Schmitz, U.: Network-Based and Machine-Learning Approaches Identify Diagnostic and Prognostic Models for EMT-Type Gastric Tumors. Genes 14(3), 750 (2023) 10.3390/genes14030750

[63] Wang, G.-C., Gan, X., Zeng, Y.-Q., Chen, X., Kang, H., Huang, S.- W., Hu, W.-H.: The Role of NCS1 in Immunotherapy and Prognosis of Human Cancer. Biomedicines 11(10), 2765 (2023) 10.3390/biomedicines11102765

[64] Xue, L., Liu, X., Wang, Q., Liu, C.Q., Chen, Y., Jia, W., Hsie, R., Chen, Y., Luh, F., Zheng, S., Yen, Y.: Ribonucleotide reductase subunit M2B deficiency leads to mitochondrial permeability transition pore opening and is associated with aggressive clinicopathologic manifestations of breast cancer. American Journal of Translational Research 10(11), 3635–3649 (2018)

[65] Zhong, X., Xiu, H., Bi, Y., Zhang, H., Chang, L., Diao, H.: Targeting eIF5A2 inhibits prostate carcinogenesis, migration, invasion and metastasis in vitro and in vivo. Bioengineered 11(1), 619–627 (2020) 10.1080/21655979.2020.1774993

[66] Xu, C., Song, L., Peng, H., Yang, Y., Liu, Y., Pei, D., Guo, J., Liu, N., Liu, J., Li, X., Li, C., Kang, Z.: Clinical Eosinophil-Associated Genes can Serve as a Reliable Predictor of Bladder Urothelial Cancer. Frontiers in Molecular Biosciences 9 (2022) 10.3389/fmolb.2022.963455

[67] Debaugny, R.E., Skok, J.A.: CTCF and CTCFL in cancer. Current Opinion in Genetics & Development 61, 44–52 (2020) 10.1016/j.gde.2020.02.021

[68] Meynet, O., Scotlandi, K., Pradelli, E., Manara, M.C., Colombo, M.P., Schmid-Antomarchi, H., Picci, P., Bernard, A., Bernard, G.: Xg expression in Ewing’s sarcoma is of prognostic value and contributes to tumor invasiveness. Cancer Research 70(9), 3730–3738 (2010) 10.1158/0008-5472.CAN-09-2837

[69] Shi, K., Shan, Y., Sun, X., Chen, K., Luo, Q., Xu, Q.: TP53INP2 modulates the malignant progression of colorectal cancer by reducing the inactive form of *β*-catenin. Biochemical and Biophysical Research Communications 690, 149275 (2024) 10.1016/j.bbrc.2023.149275

[70] Wu, I., Moses, M.A.: BNF-1, a novel gene encoding a putative extracellular matrix protein, is overexpressed in tumor tissues. Gene 311, 105–110 (2003) 10.1016/s0378-1119(03)00563-8

[71] Feng, D.-d., Cao, Q., Zhang, D.-q., Wu, X.-l., Yang, C.-x., Chen, Y.-f., Yu, T., Qi, H.-x., Zhou, G.-p.: Transcription factor E2F1 positively regulates interferon regulatory factor 5 expression in non-small cell lung cancer. OncoTargets and therapy 12, 6907–6915 (2019) 10.2147/OTT.S215701

[72] Uchida, A., Seki, N., Mizuno, K., Misono, S., Yamada, Y., Kikkawa, N., Sanada, H., Kumamoto, T., Suetsugu, T., Inoue, H.: Involvement of dual-strand of the miR-144 duplex and their targets in the pathogenesis of lung squamous cell carcinoma. Cancer Science 110(1), 420–432 (2019) 10.1111/cas.13853

[73] Cho, E.-C., Kuo, M.-L., Liu, X., Yang, L., Hsieh, Y.-C., Wang, J., Cheng, Y., Yen, Y.: Tumor suppressor FOXO3 regulates ribonucleotide reductase subunit RRM2B and impacts on survival of cancer patients. Oncotarget 5(13), 4834–4844 (2014) 10.18632/oncotarget.2044

[74] Chen, C., Zhang, B., Wu, S., Song, Y., Li, J.: Knockdown of EIF5A2 inhibits the malignant potential of non-small cell lung cancer cells. Oncology Letters 15(4), 4541–4549 (2018) 10.3892/ol.2018.7832

[75] Hong, J.A., Kang, Y., Abdullaev, Z., Flanagan, P.T., Pack, S.D., Fischette, M.R., Adnani, M.T., Loukinov, D.I., Vatolin, S., Risinger, J.I., Custer, M., Chen, G.A., Zhao, M., Nguyen, D.M., Barrett, J.C., Lobanenkov, V.V., Schrump, D.S.: Reciprocal binding of CTCF and BORIS to the NY-ESO-1 promoter coincides with derepression of this cancer-testis gene in lung cancer cells. Cancer Research 65(17), 7763–7774 (2005) 10.1158/0008-5472.CAN-05-0823

[76] Chen, H., Pan, R., Li, H., Zhang, W., Ren, C., Lu, Q., Chen, H., Zhang, X., Nie, Y.: CHRDL2 promotes osteosarcoma cell proliferation and metastasis through the BMP-9/PI3K/AKT pathway. Cell Biology International 45(3), 623–632 (2021) 10.1002/cbin.11507

[77] Sun, J., Liu, X., Gao, H., Zhang, L., Ji, Q., Wang, Z., Zhou, L., Wang, Y., Sui, H., Fan, Z., Li, Q.: Overexpression of colorectal cancer oncogene CHRDL2 predicts a poor prognosis. Oncotarget 8(7), 11489–11506 (2016) 10.18632/oncotarget.14039

[78] Lao, L., Shen, J., Tian, H., Zhong, G., Murray, S.S., Wang, J.C.: Secreted phosphoprotein 24kD (Spp24) inhibits growth of hepatocellular carcinoma in vivo. Environmental Toxicology and Pharmacology 51, 51–55 (2017) 10.1016/j.etap.2017.03.001

[79] Chen, H., Li, C., Zhou, T., Li, X., Duarte, M.E.L., Daubs, M.D., Buser, Z., Brochmann, E.J., Wang, J.C., Murray, S.S., Jiao, L., Tian, H.: Secreted phosphoprotein 24 kD (Spp24) inhibits the growth of human osteosarcoma through the BMP-2/Smad signaling pathway. Journal of Orthopaedic Research: Official Publication of the Orthopaedic Research Society 41(8), 1803–1814 (2023) 10.1002/jor.25517

[80] Lao, L., Shen, J., Tian, H., Yao, Q., Li, Y., Qian, L., Murray, S.S., Wang, J.C.: Secreted Phosphoprotein 24 kD Inhibits Growth of Human Prostate Cancer Cells Stimulated by BMP-2. Anticancer Research 36(11), 5773–5780 (2016) 10.21873/anticanres.11161

[81] Li, C.-S., Tian, H., Zou, M., Zhao, K.-W., Li, Y., Lao, L., Brochmann, E.J., Duarte, M.E.L., Daubs, M.D., Zhou, Y.-H., Murray, S.S., Wang, J.C.: Secreted phosphoprotein 24 kD (Spp24) inhibits growth of human pancreatic cancer cells caused by BMP-2. Biochemical and Biophysical Research Communications 466(2), 167–172 (2015) 10.1016/j.bbrc.2015.08.124

[82] Lee, K.-B., Murray, S.S., Duarte, M.E.L., Spitz, J.F., Johnson, J.S., Song, K.-J., Brochmann, E.J., Taghavi, C.E., Keorochana, G., Liao, J.-C., Wang, J.C.: Effects of the bone morphogenetic protein binding protein spp24 (secreted phosphoprotein 24 kD) on the growth of human lung cancer cells. Journal of Orthopaedic Research: Official Publication of the Orthopaedic Research Society 29(11), 1712–1718 (2011) 10.1002/jor.21383

[83] Langenfeld, E., Hong, C.C., Lanke, G., Langenfeld, J.: Bone Morphogenetic Protein Type I Receptor Antagonists Decrease Growth and Induce Cell Death of Lung Cancer Cell Lines. PLOS ONE 8(4), 61256 (2013) 10.1371/journal.pone.0061256

[84] Wu, C.-K., Wei, M.-T., Wu, H.-C., Wu, C.-L., Wu, C.-J., Liaw, H., Su, W.- P.: BMP2 promotes lung adenocarcinoma metastasis through BMP receptor 2-mediated SMAD1/5 activation. Scientific Reports 12(1), 16310 (2022) 10.1038/s41598-022-20788-2

[85] Zhou, W., Yan, K., Xi, Q.: BMP signaling in cancer stemness and differentiation. Cell Regeneration 12, 37 (2023) 10.1186/s13619-023-00181-8

[86] Hao, F., Zhu, Q., Lu, L., Sun, S., Huang, Y., Zhang, J., Liu, Z., Miao, Y., Jiao, X., Chen, D.: EIF5A2 Is Highly Expressed in Anaplastic Thyroid Carcinoma and Is Associated With Tumor Growth by Modulating TGF-*β* Signals. Oncology Research 28(4), 345–355 (2020) 10.3727/096504020X15834065061807

[87] Wu, R., Zhong, Q., Liu, H., Liu, S.: MicroRNA-577/EIF5A2 axis suppressed the proliferation of DDP-resistant nasopharyngeal carcinoma cells by blocking TGF- *β* signaling pathway. Chemical Biology & Drug Design 102(4), 815–827 (2023) 10.1111/cbdd.14293

[88] Zhao, G., Zhang, W., Dong, P., Watari, H., Guo, Y., Pfeffer, L.M., Tigyi, G., Yue, J.: EIF5A2 controls ovarian tumor growth and metastasis by promoting epithelial to mesenchymal transition via the TGF*β* pathway. Cell & Bioscience 11(1), 70 (2021) 10.1186/s13578-021-00578-5

[89] Zhang, Z., He, G., Lv, Y., Liu, Y., Niu, Z., Feng, Q., Hu, R., Xu, J.: HERC3 regulates epithelial-mesenchymal transition by directly ubiquitination degradation EIF5A2 and inhibits metastasis of colorectal cancer. Cell Death & Disease 13(1), 1–12 (2022) 10.1038/s41419-022-04511-7

[90] Zhou, Z., Liu, X., Li, Y., Li, J., Deng, W., Zhong, J., Chen, L., Li, Y., Zeng, X., Wang, G., Zhu, J., Fu, B.: TP53INP2 Modulates Epithelial-to-Mesenchymal Transition via the GSK-3*β*/*β*-Catenin/Snail1 Pathway in Bladder Cancer Cells. OncoTargets and therapy 13, 9587–9597 (2020) 10.2147/OTT.S251830

[91] Soltanian, S., Dehghani, H.: BORIS: A key regulator of cancer stemness. Cancer Cell International 18, 154 (2018) 10.1186/s12935-018-0650-8

[92] Janssen, S.M., Moscona, R., Elchebly, M., Papadakis, A.I., Redpath, M., Wang, H., Rubin, E., van Kempen, L.C., Spatz, A.: BORIS/CTCFL promotes a switch from a proliferative towards an invasive phenotype in melanoma cells. Cell Death Discovery 6, 1 (2020) 10.1038/s41420-019-0235-x

[93] Makani, V.K.K., Mendonza, J.J., Edathara, P.M., Yerramsetty, S., Pal Bhadra, M.: BORIS/CTCFL expression activates the TGF*β* signaling cascade and induces Drp1 mediated mitochondrial fission in neuroblastoma. Free Radical Biology & Medicine 176, 62–72 (2021) 10.1016/j.freeradbiomed.2021.09.010

[94] De Marzio, M., Glass, K., Kuijjer, M.L.: Single-sample network modeling on omics data. BMC Biology 21(1), 296 (2023) 10.1186/s12915023-01783-z

